# The allosteric control mechanism of bacterial glycogen biosynthesis disclosed by cryoEM

**DOI:** 10.1101/2020.01.24.916759

**Authors:** Javier O. Cifuente, Natalia Comino, Cecilia D’Angelo, Alberto Marina, David Gil-Carton, David Albesa-Jové, Marcelo E. Guerin

## Abstract

Glycogen and starch are the major carbon and energy reserve polysaccharides in nature, providing living organisms with a survival advantage. The evolution of the enzymatic machinery responsible for the biosynthesis and degradation of such polysaccharides, led the development of mechanisms to control the assembly and disassembly rate, to store and recover glucose according to cell energy demands. The tetrameric enzyme ADP-glucose pyrophosphorylase (AGPase) catalyzes and regulates the initial step in the biosynthesis of both α-polyglucans. Importantly, AGPase displays cooperativity and allosteric regulation by sensing metabolites from the cell energy flux. The understanding of the allosteric signal transduction mechanisms in AGPase arises as a long-standing challenge. In this work, we disclose the cryoEM structures of the paradigmatic homotetrameric AGPase from *Escherichia coli* (*Ec*AGPase), in complex with either positive or negative physiological allosteric regulators, FBP and AMP respectively, both at 3.0 Å resolution. Strikingly, the structures reveal that FBP binds deeply into the allosteric cleft and overlaps the AMP site. As a consequence, FBP promotes a concerted conformational switch of a regulatory loop, RL2, from a ‘locked’ to a ‘free’ state, modulating ATP binding and activating the enzyme. This notion is strongly supported by our complementary biophysical and bioinformatics evidence, and a careful analysis of vast enzyme kinetics data on single-point mutants of *Ec*AGPase. The cryoEM structures uncover the residue interaction networks (RIN) between the allosteric and the catalytic components of the enzyme, providing unique details on how the signaling information is transmitted across the tetramer, from which cooperativity emerges. Altogether, the conformational states visualized by cryoEM reveal the regulatory mechanism of *Ec*AGPase, laying the foundations to understand the allosteric control of bacterial glycogen biosynthesis at the molecular level of detail.

## INTRODUCTION

Glucose is a central carbon and energy currency in nature. The evolution leaded organisms to acquire specific pathways to store glucose in the form of α-glucan polysaccharides. The animal and bacterial glycogen, and the plant starch are the classical functional glucose storages in the cell.^1^ In addition, α-glucans were found as essential structural components of the bacterial capsule and biofilms playing an important role in pathogenesis.^2-4^ Glycogen and starch are large homopolymers composed by linear chains of α-(1→4)-glucose residues, containing α-(1→6)-linkages at branching points (Figure 1A).^1,5^ These chemical structures provide a high number of reactive-ends that facilitates rapid storage and recovery of glucose.^6^ Glucose activation is the first step required to overcome the energy barrier for subsequent polymerization (Figure 1B). The classical pathway for bacterial glycogen biosynthesis involves the action of three enzymes: ADP-glucose pyrophosphorylase (AGPase), glycogen synthase (GS) and glycogen branching enzyme (GBE; Figure 1B).^5,7-9^ AGPase catalyzes the biosynthesis of the activated-sugar ADP-glucose (Figures 1B-C), whereas GS generates a linear α-(1→4)-linked glucose chain, and the GBE produces α-(1→6)-linked glucan branches in the polymer. ^5,7-9^ Glycogen degradation is carried out by glycogen phosphorylase (GP), which functions as a depolymerizing enzyme, and the glycogen debranching enzyme (GDE) that catalyzes the removal of α-(1→6)-linked ramifications (Figure 1B).^5^

**Figure 1.**
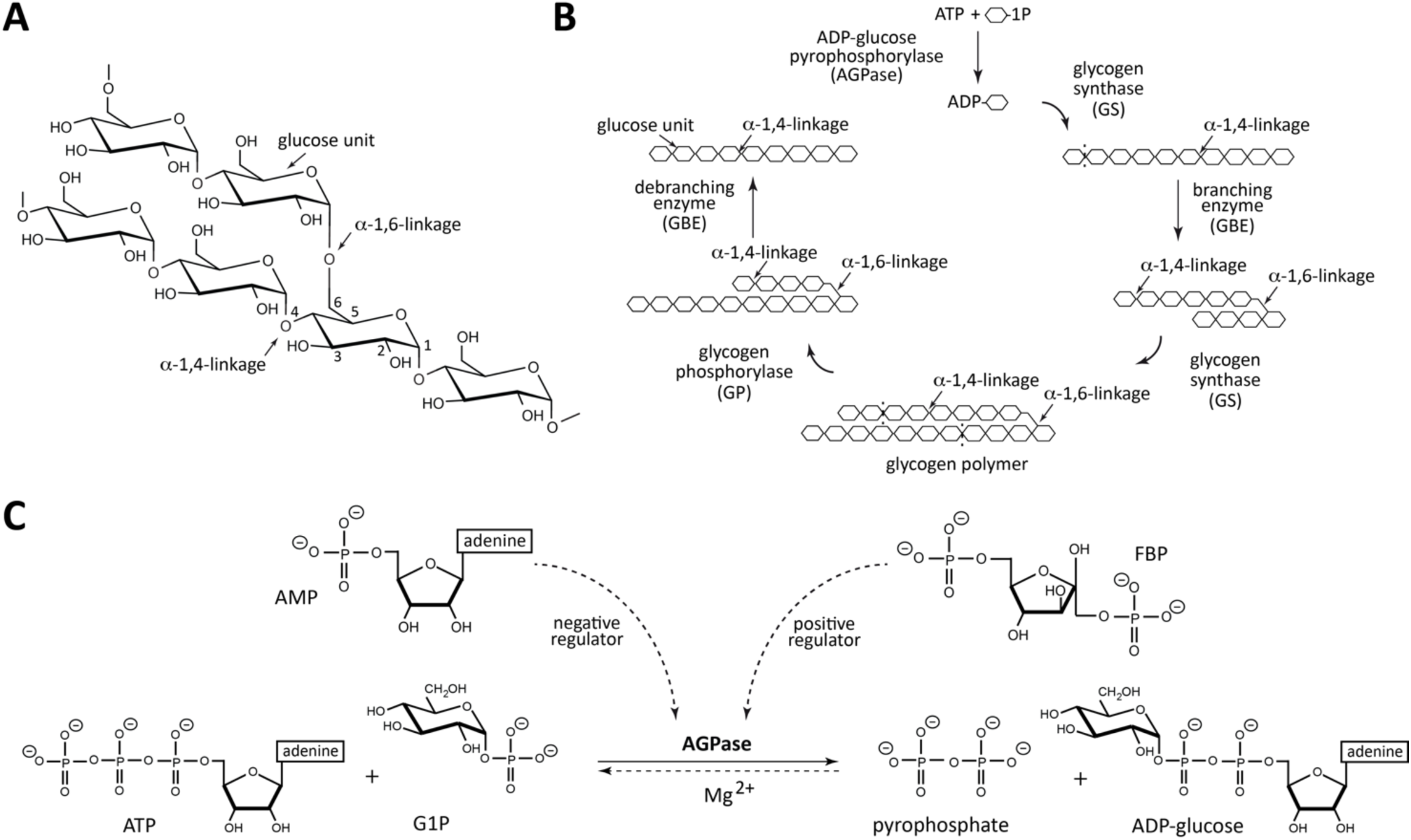
Classical bacterial glycogen structure, biosynthesis and regulation. (**A**) Glycogen is a very large-branched glucose homopolymer containing ∼90% α-(1→4)-glucosidic linkages and 10% α-(1→6)-linkages. (**B**) The classical bacterial glycogen biosynthetic pathway involves the action of three enzymes: AGPase, GS; and GBE. Glycogen degradation is carried out by two enzymes: GP and GDE. (**C**) *Ec*AGPase catalyzes the main regulatory step in bacterial glycogen. *Ec*AGPase catalyzes the reaction between ATP and G1P in the presence of a divalent metal cation, Mg^2+^, to form ADP-Glc and PPi. ADP-Glc biosynthesis and hydrolysis directions are shown as a full and dotted lines, respectively. The two major positive and negative allosteric regulators, FBP and AMP, respectively, are shown.

Cell homeostasis requires precise and rapid mechanisms to sense the organism status to coordinate the metabolic network accordingly.^10^ To oversight the cell energy storage, glycogen and starch biosynthetic pathways are controlled by AGPase through cooperative and allosteric strategies.^11^ The cooperativity emerges when the interaction of one molecule modulates the binding of another molecule of the same nature in other protomers, while allosterism arises when the association of a molecule modulates the binding of a different type of molecule to the oligomeric architecture of the enzyme.^11-14^ AGPase catalyzes the reversible condensation reaction between adenosine triphosphate (ATP) and glucose 1-phosphate (G1P) to produce ADP-glucose and pyrophosphate (PPi; Figure 1C).^5,15,16,17^ Specifically, the oxygen on the phosphate group of G1P acts as a nucleophile attacking the α-PO_4_ group of the nucleoside triphosphate, leading to the liberation of PPi (Figure S1).^18^ The reaction is held in the presence of the divalent metal cation Mg^2+^, which minimizes the charge repulsion between phosphate groups, favoring nucleophile activation.^19,20^ Moreover, two positively charged residues polarize these groups, increasing the nucleophilic nature of the oxygen attacking the phosphor atom.^21,22^ AGPase follows a sequential ordered bi–bi mechanism with ATP binding first, followed by G1P and by the ordered release of PPi and ADP-glucose.^23^ The hydrolysis of PPi by inorganic pyrophosphatases results in a global irreversible and energetically expensive reaction *in vivo*.^24,25^ Thus, evolution led AGPase to acquire an exquisite allosteric regulation mechanism to control its enzymatic activity by essential metabolites in the energetic flux within the cell.^5,15,17^ AGPase activators are metabolites that reflect signals of high carbon and energy content of a particular bacteria or tissue, whereas inhibitors indicate low metabolic energy levels. As a consequence, AGPases have been classified into nine different classes taking into account the specific positive or negative allosteric regulators.^17^

Bacterial AGPases are encoded by a single gene, giving rise to a native homotetrameric protein (α4) with a molecular mass of ca. 200 kDa (ca. 50 kDa for each protomer).^5,17^ In contrast, plant AGPases are composed of two α and two β subunits, also referred to as ‘small’ and ‘large’ subunits, respectively, to form an α2β2 heterotetramer.^26,27,28,29^ The two subunits have different functions; α is the catalytic subunit, whereas β is the regulatory subunit. To date, three crystal structures of AGPases have been reported, those of the bacterial homotetrameric AGPases from *Escherichia coli* (*Ec*AGPase)^5,16,30^ and *Agrobacterium tumefaciens* (*At*AGPase),^31,32^ and a recombinant homotetrameric version of the small subunit (α4) of the photosynthetic potato tuber AGPase (*St*AGPase).^33^ The activity of the paradigmatic *Ec*AGPase is enhanced by the high-energy glycolytic intermediate fructose-1,6-bisphosphate (FBP), whereas, the ubiquitous low-energy metabolite AMP inhibits the enzyme. Thus, the interplay of both regulators tune *Ec*AGPase activity.^5,15,34^ Moreover, *Ec*AGPase show positive cooperativity among ATP sites and a considerable *K*_M_ reduction by FBP.^34-37^ We reported the first crystal structures of the paradigmatic *Ec*AGPase in complex with FBP and AMP, respectively.^16,30^ These structures provided a first perspective of the regulatory sites and the identification of secondary structure elements likely participating into the allosteric regulation. However, the structural data prohibited the visualization of the architecture of *Ec*AGPase in biologically relevant conformations and the comprehension on how the allosteric regulators act in a concerted manner to control the enzymatic activity. The structural framework that modulates these properties was historically provided by x-ray crystallography studies. However, the organization of the protein molecules inside the crystal packing prevents, in many cases, the visualization of native biologically relevant conformational states associated to the binding of substrates and regulatory molecules, impeding the construction of accurate models to explain allosteric regulation.^16,31-33^ In this work, we disclose the cryoEM structures of the homotetrameric *Ec*AGPase (a ca. 200 kDa enzyme, with each protomer of 48.7 kDa), in complex with FBP and AMP, both at 3.0 Å resolution, respectively. In combination with biochemical, biophysical and bioinformatics data, the regulatory and cooperativity mechanisms of *Ec*AGPase is unraveled, laying the foundations to understand the allosteric control of bacterial glycogen biosynthesis.

## RESULTS

### The *Ec*AGPase-FBP complex as visualized by cryoEM

To understand how the positive and negative signals are propagated across the *Ec*AGPase structure in the native state in solution, we determined the single-particle cryoEM structures of *Ec*AGPase in complex with either regulator, FBP or AMP (Figures 2 and 3; Figures S2 to S7; Table 1). *Ec*AGPase is a homotetrameric enzyme that can be visualized as a dimer-of-dimers, with each protomer comprising an N-terminal glycosyltransferase A-like (GT-A-like) catalytic domain, and a C-terminal Left-Handed-β-helix (LβH) regulatory domain (Figures 2 and 3; Figure S6; Video S1: https://www.dropbox.com/s/fupmbncoxj9yo5k/vf_Video_S1.avi?dl=0).^16^ The three-dimensional reconstruction of the *Ec*AGPase-FBP complex was carried out without imposing symmetry operators, C1 (*Ec*AGPase-FBP_C1_; 3.24 Å resolution), enforcing C2 symmetry (*Ec*AGPase-FBP_C2_; 3.16 Å resolution), and imposing D2 symmetry (*Ec*AGPase-FBP_D2_; 3.05 Å resolution) (Figures S2, S4 and S6-S7). The similar resolution achieved by these reconstructions and the high correlation among them explain the highly symmetrical characteristic of this complex. The *Ec*AGPase-FBP_D2_ reconstruction reveals a consensus density for FBP among the four regulatory clefts. Strikingly, this binding mode shows a different location of the positive regulator compared to that observed in the *Ec*AGPase-FBP complex obtained by X-ray crystallography (Figures 2B-C and 4A-B).^16,38^ The FBP is found deeply buried into the cleft mainly defined by (i) the N-terminal β2-β3 hairpin (residues 46–52), α5, and the connecting loop α2-α3 (residues 37–42), which partially constitutes the so call ‘sensory motif’ (SM), communicating the regulatory and active sites of each subunit, and (ii) the C-terminal α15 (residues 419–425) and the connecting loops β28-β29 (residues 384–388) and β25-β26 (residues 367–371; Figures 2A-C). The first PO_4_ group of FBP occupies a cavity rich in positively charged residues including the side chains of Arg40 (α3), His46 and Arg52 (β2-β3 hairpin) and Arg386 (LβH; Figures 2A-C). The O4 of the fructose ring makes a hydrogen bond with the guanidinium group of Arg419. The second PO_4_ group makes a critical interaction with the side chain of Lys39, essential for FBP binding and the activation of *Ec*AGPase and with the guanidinium group of Arg423.^39-40^ The positive regulator was observed in the same location in both *Ec*AGPase-FBP_C1_ and *Ec*AGPase-FBP_C2_ reconstructions. Nevertheless, minor differences in the FBP_C1_ and FBP_C2_ conformation are revealed by the subtle movements of the second PO_4_ group. Importantly, the *Ec*AGPase-FBP_C1_ reconstruction exhibits clear electron density to model only two FBP molecules in two regulatory sites located in the same dimer, although residual electron density can be observed in the other two allosteric sites, suggesting that FBP may adopt distinct local conformations.

**Figure 2.**
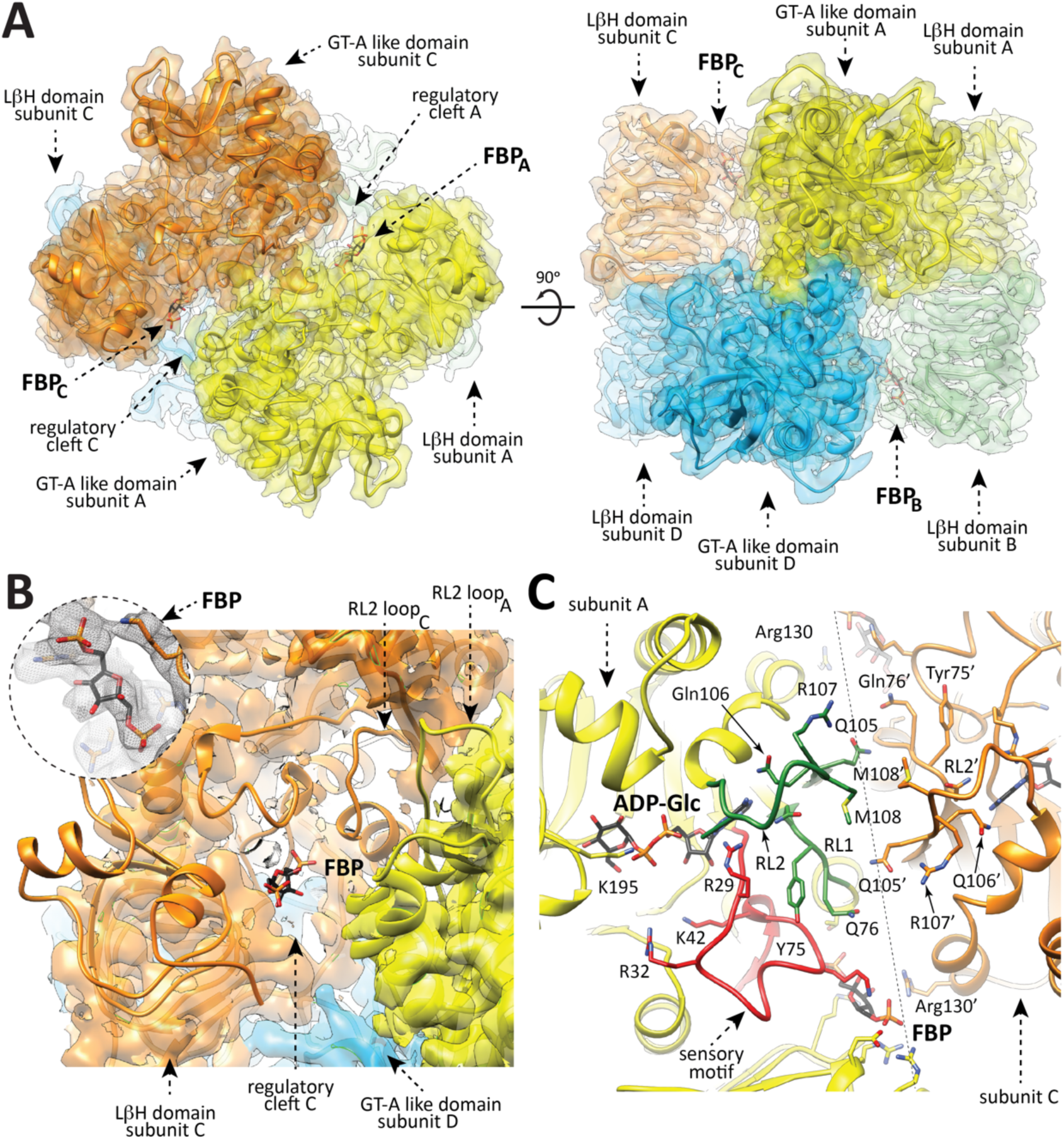
Overall structure of the *Ec*AGPase-FBP complex as visualized by cryoEM. (**A**) The overall view of the electron density cryoEM map of the *Ec*AGPase-FBP_D2_ complex, colored according to the four protomers (orange, yellow, blue and green) of the homotetramer. (**B**) A selected region corresponding to the neighborhood of the FBP binding site is masked to reveal the atomic model built into the electron density, with the protein shown as ribbons and the ligand FBP in sticks representation. Close-up of the *Ec*AGPase-FBP_D2_ regulatory site showing the location and the electron density cryoEM map of the activator FBP. (**C**) Cartoon representation of the interface between the reference protomer (yellow) and the neighbor subunit (orange) of the *Ec*AGPase-FBP_D2_ reconstruction. Note that the RL2 loop conformation as observed in the *Ec*AGPase-FBP_D2_ reconstruction faces its own active site.

**Figure 3.**
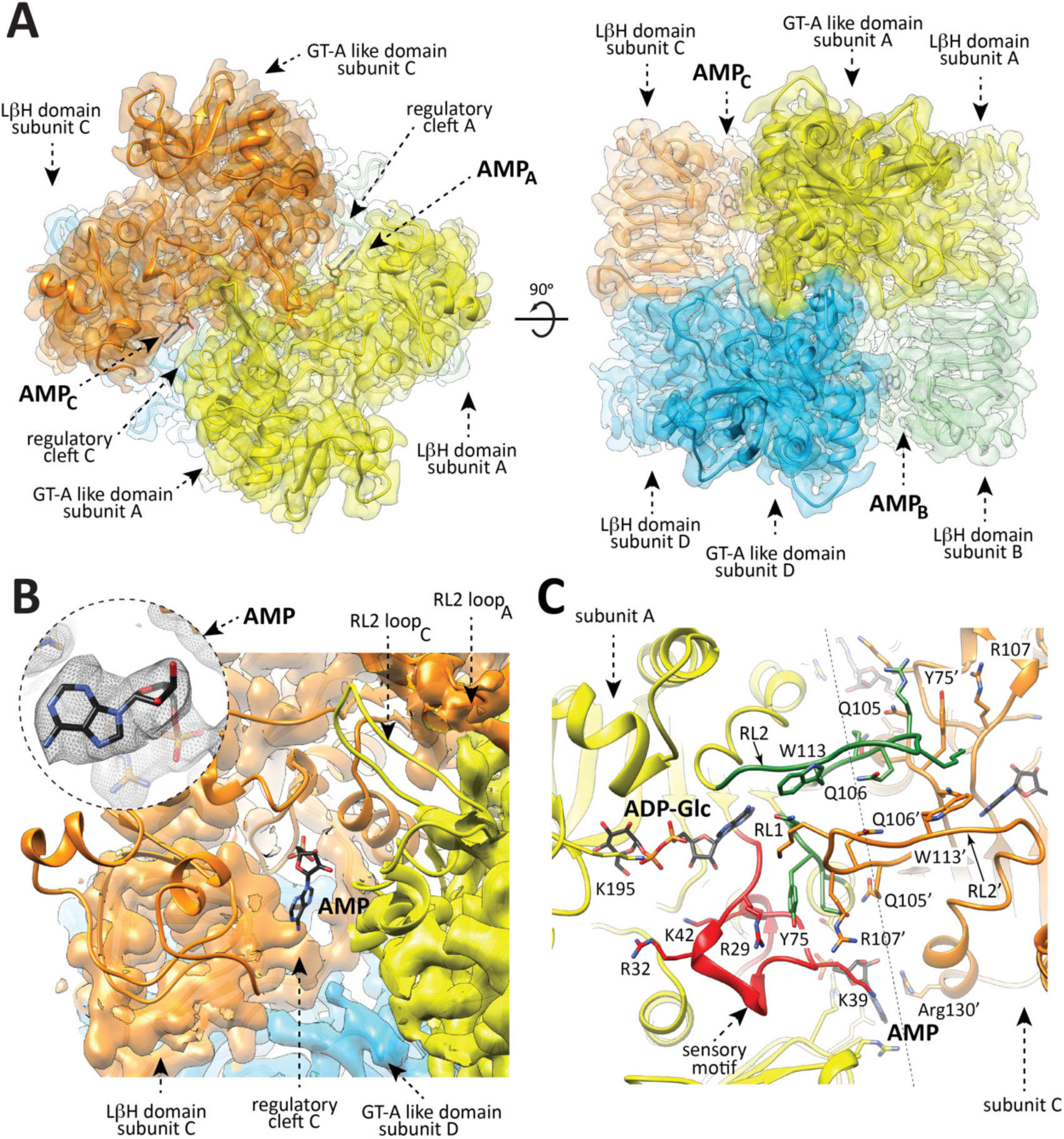
Overall structure of the *Ec*AGPase-AMP complex as visualized by cryoEM. (**A**) The overall view of the electron density cryoEM map of the *Ec*AGPase-AMP_D2_ complex, colored according to the four protomers (orange, yellow, blue and green) of the homotetramer. (**B**) A selected region corresponding to the neighborhood of the AMP binding site is masked to reveal the atomic model built into the electron density, with the protein shown as ribbons and the ligand AMP in sticks representation. Close-up of the *Ec*AGPase-AMP_D2_ regulatory site showing the location and the electron density cryoEM map of the activator AMP. (**C**) Cartoon representation of the interface between the reference protomer (yellow) and the neighbor subunit (orange) of the *Ec*AGPase-AMP_D2_ reconstruction. Note that the RL2 loop conformation, as observed in the *Ec*AGPase-AMP_D2_ reconstruction, is oriented towards the neighbor active site.

**Figure 4.**
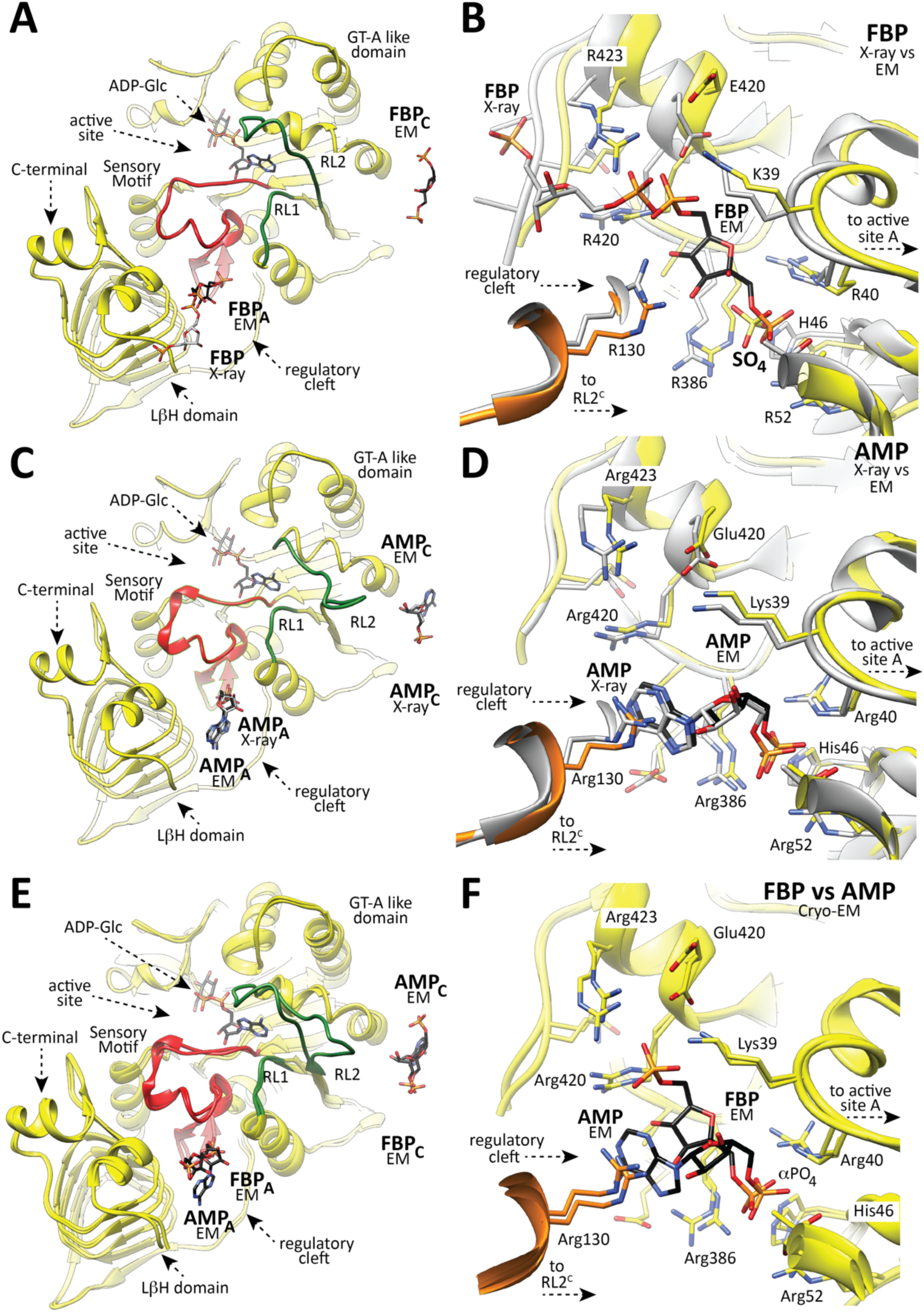
The location of FBP and AMP regulators into the allosteric site as observed by cryoEM. (**A**) Cartoon representation of one protomer of *Ec*AGPase as observed in the *Ec*AGPase-FBP_D2_ complex obtained by cryoEM. The secondary structure elements involved in the regulatory mechanism of *Ec*AGPase are shown: (i) the SM (red) communicates the regulatory cleft and active sites of the same protomer, (ii) the RL1 (green) and (iii) RL2 loops (green). The FBP molecule is shown in black. The location of the FBP molecule as observed in the X-ray *Ec*AGPase-FBP complex is shown in grey as a reference. (**B**) Closed view of the regulatory site as observed in the cryoEM *Ec*AGPase-FBP_D2_ complex (yellow) and the *Ec*AGPase-FBP complex (grey) obtained by X-ray crystallography. The FBP molecule is shown in black and grey, respectively. A molecule of ADP-Glc is shown in the active site as a reference. Note that the sulfate ion occupying the cleft cavity belongs to the structure of the *Ec*AGPase-FBP complex solved by X-ray crystallography. (**C**) Cartoon representation of one protomer of *Ec*AGPase as observed in the *Ec*AGPase-AMP_D2_ complex obtained by cryoEM, and the location of AMP (black) into the regulatory site. The location of AMP in the *Ec*AGPase-AMP complex solved by X-ray crystallography is depicted in grey. A molecule of ADP-Glc is shown in the active site as a reference. (**D**) Closed view of the regulatory site as observed in the *Ec*AGPase-AMP complex obtained by cryoEM (*Ec*AGPase-AMP_D2_; yellow) and X-ray crystallography (grey). The AMP molecule is shown in black. The location of the AMP molecule as observed in the X-ray *Ec*AGPase-AMP complex is shown in grey as a reference. (**E**) Structural superposition of one protomer in the *Ec*AGPase-FBP_D2_ and *Ec*AGPase-AMP_D2_ complexes obtained by cryoEM. The FBP and AMP regulators are shown. A molecule of ADP-Glc is shown in the active site as a reference. (**F**) Closed view of the regulatory site as observed in the *Ec*AGPase-FBP_D2_ and *Ec*AGPase-AMP_D2_ complexes.

**Figure 5.**
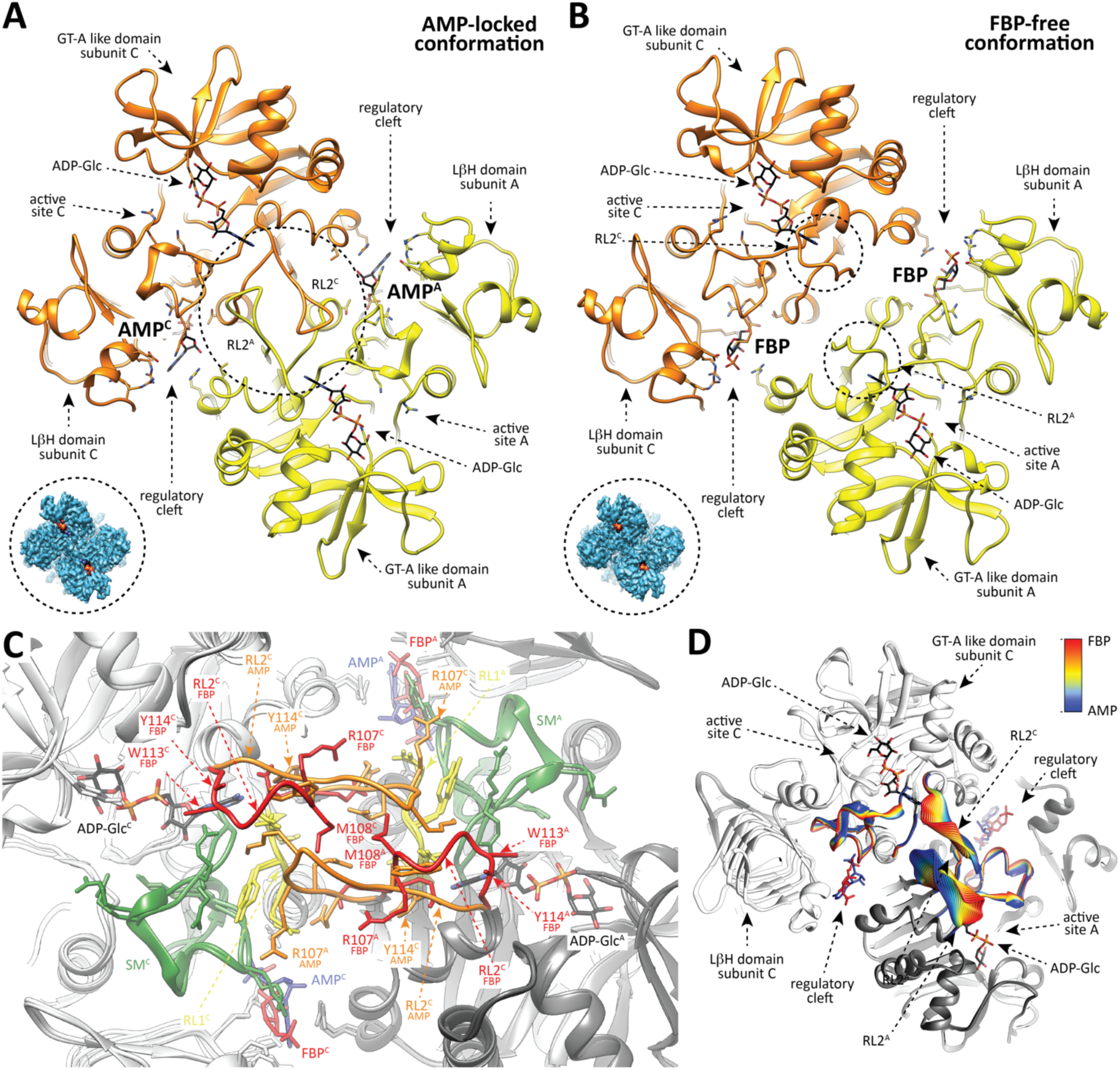
The conformation of the RL2 loop as observed in the *Ec*AGPase-FBP_D2_ and *Ec*AGPase-AMP_D2_ complexes. (**A**) View of two neighboring protomers of different dimers (yellow and orange) in the *Ec*AGPase-FBP_D2_ complex. FBP is observed in the regulatory cleft, whereas ADP-Glc is shown in the active site as a reference. The “free” conformation of the RL2 loops is indicated. A caption of the *Ec*AGPase-FBP_D2_ complex map (blue) is shown with ADP-Glc placed in the active site. (**B**) As in panel (A), two neighboring protomers of different dimers in the *Ec*AGPase-AMP_D2_ complex. AMP is observed in the regulatory cleft, whereas ADP-Glc is shown in the active site as a reference. The “locked” conformation of the RL2 loops is indicated. A caption of the *Ec*AGPase-AMP_D2_ complex map (blue) is shown with ADP-Glc to observe map differences of the active shape compared with the *Ec*AGPase-FBP_D2_ complex. (**C**) Structural comparison of the RL1 and RL2 loops as observed in the *Ec*AGPase-FBP_D2_ and *Ec*AGPase-AMP_D2_ complexes. (**D**) Cartoon representation showing the transition of the SM, RL1 and RL2 loops conformation as observed by cryoEM. Different positions of the main chain are colored according to the color scheme shown in the bar (FBP complex red, AMP complex blue).

**Figure 6.**
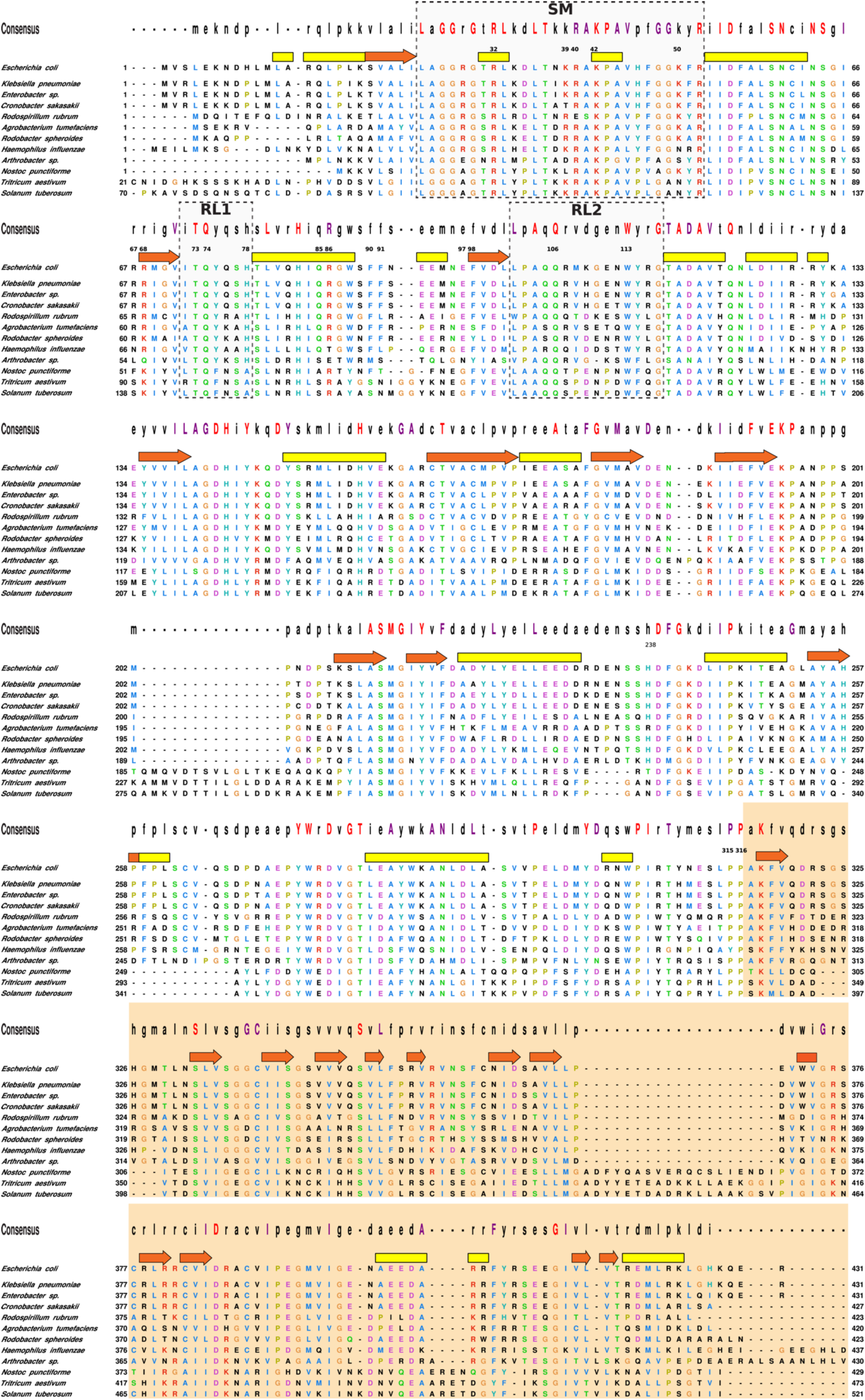
Structural weighted alignment of *Ec*AGPase with selected homologs. One-letter code residues are shown colored according to Clustal X scheme. The consensus sequence is displayed on top of the alignment where highly conserved residues in capitalized one-letter code (completely conserved residues in red, purple colored conservation ≥ 80%). The white background indicates the region corresponding to the N-terminal catalytic domain meanwhile the light orange background agrees with the C-terminal domain. Yellow and red bars indicate either α-helices and β-strand regions comprised in the *Ec*AGPase structure, respectively. Regions corresponding to the sensory motif (SM), and the regulatory loops (RL1 and RL2) are enclosed in black boxes. The numbering of relevant *Ec*AGPase residues is indicated. Shown are AGPases from *E. coli* (UniProt code P0A6V1), *Klebsiella pneumoniae* (B5XTQ9), *Enterobacter sp.* (A4WFL3), *Cronobacter sakasakii* (A7MGF4), *Rodospirillum rubrum* (Q2RS49), *Agrobacterium tumefaciens* (P39669), *Rodobacter spheroids* (A3PJX6), *Haemofilus influenza* (P43796), *Arthrobacter sp.* (A0JWV0), *Nostoc punctiforme* (B2IUY3), *Triticum aestivum* (P30523), and *Solanum tuberosum* (P23509).

**Figure 7.**
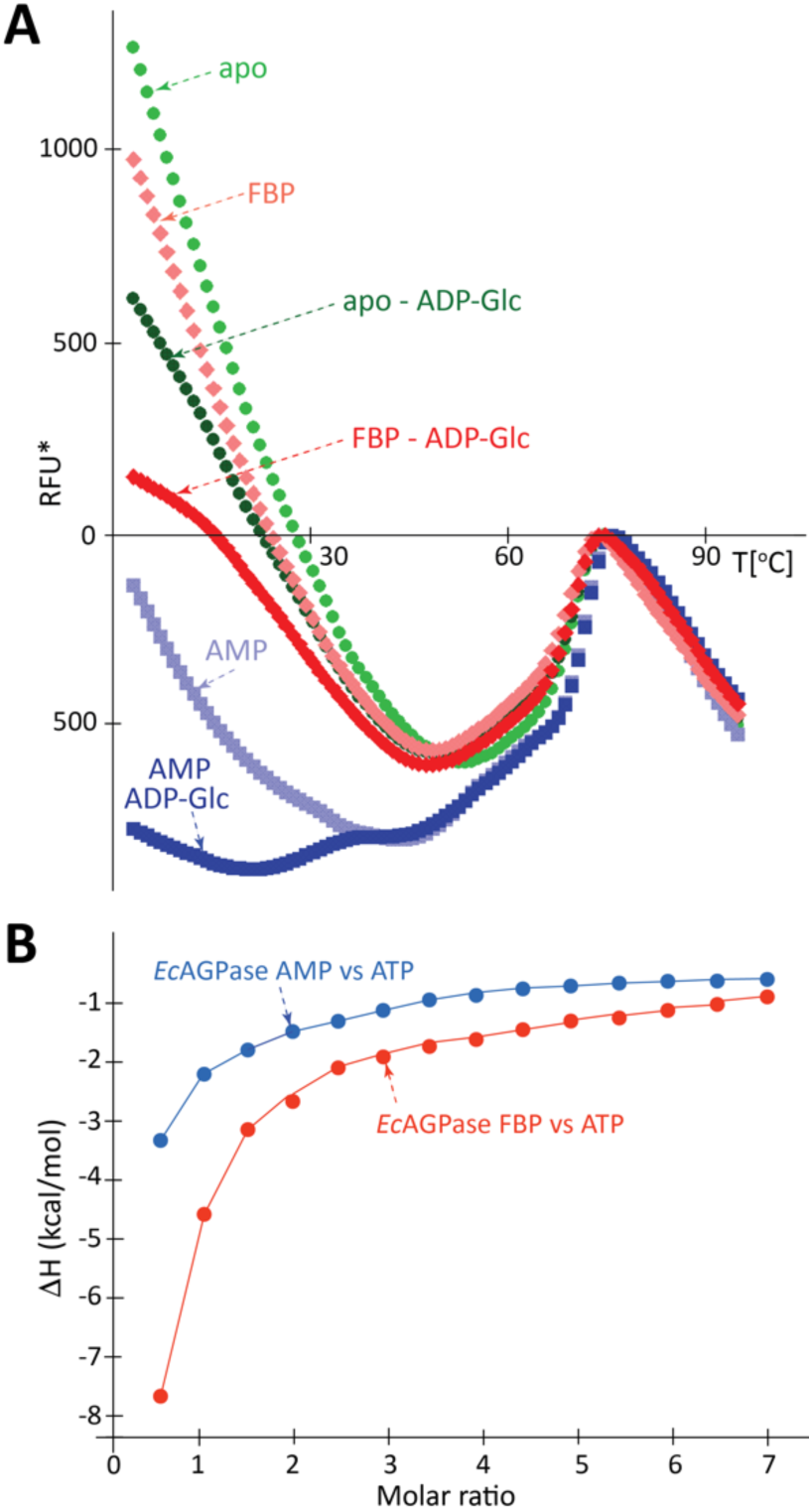
The allosteric regulators modulate the affinity for the substrate ATP. (**A**) Accessibility of ATP to the *Ec*AGPase active site is modulated by the presence of the allosteric regulators FBP and AMP. (**B**) ITC measurements of *Ec*AGPase-ligand interactions. The integrated heats of injections of the titrations corrected for the ligand heat of dilution and normalized to the ligand concentration. Solid lines correspond to the best fit of data.

**Table 1.** CryoEM data collection, single-particle reconstruction maps, and model statistics.

| Data collection information |  |  |  |  |  |  |
| --- | --- | --- | --- | --- | --- | --- |
| Data set | *EcAGPase*-FBP complex |  |  | *EcAGPase*-AMP complex |  |  |
| Nominal magnification | 47755x |  |  | 47755x |  |  |
| Voltage (kV) | 300 |  |  | 300 |  |  |
| Electron dose (e-/Å<sup>2</sup>) | 45.6 |  |  | 44.8 |  |  |
| Pixel size (Å) | 1.047 |  |  | 1.047 |  |  |
| Movie number | 2975 |  |  | 1681 |  |  |
| Defocus average and range (µm) | 1.4<br>(0.2 - 2.6) |  |  | 1.7<br>(0.3 - 5.0) |  |  |
| Frames | 40 |  |  | 40 |  |  |
| Single-particle reconstruction information |  |  |  |  |  |  |
| Map | *EcAGPase*-FBP complex |  |  | *EcAGPase*-AMP complex |  |  |
| Final number of particle images in map | 297275 |  |  | 94769 |  |  |
| Symmetry imposed | D2 | C2 | C1 | D2 | C2 | C1 |
| EMDB code | EMD-4754 | EMD-10199 | EMD-10201 | EMD-4761 | EMD-10203 | EMD-10208 |
| Map resolution (Å)<br>FSC=0.143 criterion | 3.05 | 3.16 | 3.24 | 2.95 | 3.09 | 3.26 |
| Map sharpening<br>B factor (Å<sup>2</sup>) | -148.1 | -139.9 | 131.2 | -116.8 | -114.5 | -99.9 |
| Model information |  |  |  |  |  |  |
| Model | *EcAGPase*-FBP complex |  |  | *EcAGPase*-AMP complex |  |  |
| PDB ID | 6R8B | 6SHJ | 6SHN | 6R8U | 6SHQ | 6SI8 |
| Atoms (Non-H) | 26092 | 25974 | 25359 | 26760 | 26874 | 26590 |
| Protein residues | 1688 | 1664 | 1644 | 1700 | 1696 | 1700 |
| Ligands | 4 | 4 | 2 | 4 | 4 | 4 |
| Bonds (RMSD) |  |  |  |  |  |  |
| Length (Å) | 0.01 | 0.01 | 0.016 | 0.01 | 0.01 | 0.01 |
| Angles (°) | 1.34 | 1.01 | 1.34 | 1.35 | 1.38 | 1.40 |
| Mean B-factor |  |  |  |  |  |  |
| Protein | 153.64 | 163.46 | 94.72 | 134.96 | 136.07 | 88.79 |
| Ligand | 176.65 | 191.65 | 101.34 | 127.23 | 126.19 | 82.53 |
| MolProbity score | 2.11 | 2.02 | 1.89 | 2.30 | 2.04 | 1.79 |
| Clashscore | 7.93 | 3.54 | 2.21 | 7.17 | 5.36 | 2.14 |
| Rotamer outliers (%) | 3.12 | 3.58 | 4.81 | 3.32 | 2.28 | 2.45 |
| Ramachandran plot |  |  |  |  |  |  |
| Favored (%) | 95.71 | 92.90 | 94.47 | 91.43 | 92.06 | 91.31 |
| Allowed (%) | 4.05 | 6.98 | 5.41 | 8.57 | 7.94 | 8.33 |
| CC (volume) | 0.77 | 0.75 | 0.80 | 0.80 | 0.78 | 0.79 |
| Mean CC for ligands | 0.74 | 0.66 | 0.74 | 0.84 | 0.82 | 0.84 |
| CC (mask) | 0.77 | 0.75 | 0.81 | 0.80 | 0.79 | 0.79 |

A detailed comparison of the overall *Ec*AGPase-FBP_C1_, *Ec*AGPase-FBP_C2_ and *Ec*AGPase-FBP_D2_ reconstructions reveal remarkable structural differences in the conformation of two loops: one located at the center of the particle, hereafter called the protomer ‘core loop’ (CL; residues 88-97), and the long ‘regulatory loop’ RL2 (residues Ala104 to Gly116).^16^ Specifically, the side chains of residues Phe90 and Phe91, located in the CL loop, interact with each other in the *Ec*AGPase-FBP_C2_ and *Ec*AGPase-FBP_D2_ reconstructions. Nevertheless, in the *Ec*AGPase-FBP_C1_ reconstruction, the aromatic ring of Phe91 moves away from the GT-A like core ∼180° and interacts with Tyr309 of the neighbor dimer. This structural rearrangement results in distinct pairs of CL conformations for each dimer in the *Ec*AGPase-FBP_C1_ structure. The *Ec*AGPase-FBP_C1_ and *Ec*AGPase-FBP_C2_ reconstructions display bulky density impairing to model the position of the RL2 loop located in the ATP binding site.^16^ This electron density likely represents the RL2 loop in multiple conformational states, as suggested in our previous crystallographic studies on *Ec*AGPase, as well as in other homologues, where the it could not be modeled except when participating in crystal contacts.^16,31-33^ Conversely, the *Ec*AGPase-FBP_D2_ reconstruction reveals a consensus electron density for the RL2 loop in a relaxed ‘free’ conformation (Figures 2 and 5). Moreover, the RL1 (residues Ile72 to His78) and the G-rich loops (residues 26 to 32; a constituent of the SM) appear in different conformations in these reconstructions, revealing the underlying dynamics of these loops. These findings suggest that the *Ec*AGPase-FBP complex is a quasyD2 structure with local details that can be better recovered with C1 and C2 symmetries. Taken together, the conformations of the RL2 loop in an ‘free’ state, and that of the SM G-rich loop and the RL1 loop, represent a configuration available for ATP binding.

### The *Ec*AGPase-AMP complex as visualized by cryoEM

The *Ec*AGPase-AMP complex reconstructions also reach high resolution in all symmetries, C1 (*Ec*AGPase-AMP_C1_; 3.26 Å resolution), C2 (*Ec*AGPase-AMP_C2_; 3.09 Å resolution), and D2 (*Ec*AGPase-AMP_D2_; 2.95 Å resolution), revealing a good correlation among them (Figures 3 and 4C-D; Figures S3, S5, S6-S8; Table 1). These three reconstructions exhibit unambiguous electron density for the allosteric inhibitor in all four allosteric clefts, exhibiting an AMP-binding mode analogous to the one visualized in the *Ec*AGPase-AMP X-ray crystal structure (Figure 4D).^16^ The AMP αPO_4_ group is located deep into the positively charged cavity and coordinated by the side chains of residues Arg40, His46, Arg53 (SM), Thr79 (α5), and Arg386 (LβH). The O2 of the ribose ring makes a strong interaction with the guanidinium group of Arg130. The adenine heterocycle is also stabilized by a strong stacking interaction with Arg130 (α7) from the GT-A-like domain of a neighboring subunit. Strikingly, the *Ec*AGPase-AMP_D2_ symmetry reconstruction reveals an extraordinary structural rearrangement of the RL2 loop, also observed in the lower symmetries (Figures 2F and 3B). The RL2 loop cross-over towards the active site of a neighbor protomer from a different dimer, adopting a ‘locked’ state stabilized by important interactions with the RL2’ loop, as well as with both RL1 and RL1’ loops. In addition, the CL loop is observed in the same conformation in all applied symmetries, indicating that AMP stabilizes a highly symmetrical quaternary conformation. We previously assigned to the AMP stacking interactions the main reason for a substantial stabilization, in more than 5°C, of the enzyme.^16,30^ The cryoEM *Ec*AGPase-AMP structure discloses now the contribution of the loops in the stabilization effect.

**Figure 8.**
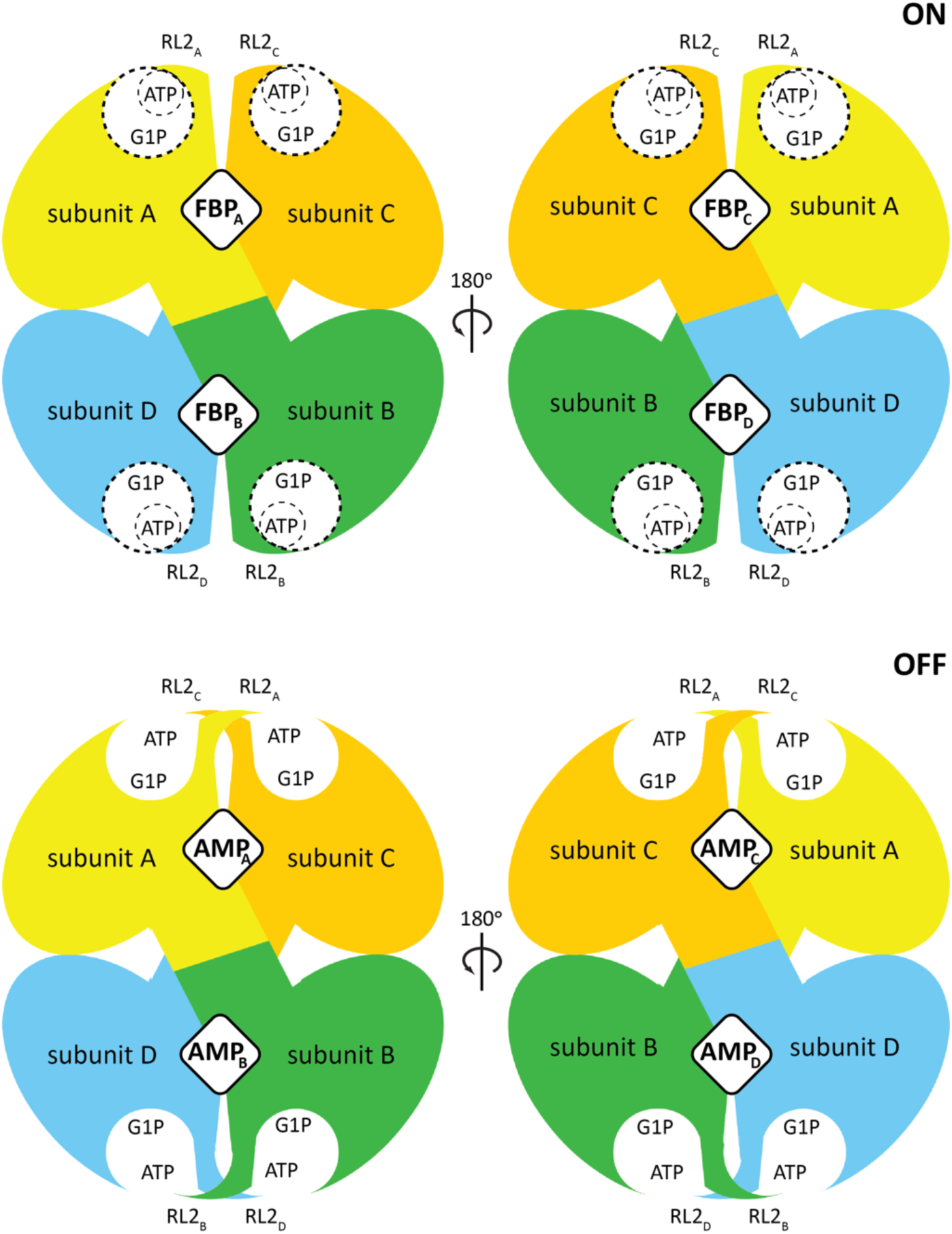
A molecular model for the allosteric regulation of *Ec*AGPase. The *Ec*AGPase tetramer is displayed in overlapping purple spheres, where the dotted lines are inter-protomer interphases. The active site is displayed as an open circle containing the substrates ATP and G1P. The allosteric clefts are indicated with rhombi at the interphases, that can alternatively be occupied by FBP or AMP, leading to the ON or OFF states, respectively. In the enzyme ON-state, the active site’s RL2 loops (depicted in colors) are in a putative conformation allowing the interaction with ATP. In contrast, in the OFF inhibited AMP-state, the RL2 loops from neighbor protomers are engaged in the “locked” state, sequestering the loops for the interaction with ATP.

The structural comparison of the *Ec*AGPase-AMP_D2_ and *Ec*AGPase-FBP_D2_ reconstructions revealed important conformational changes (Figures 4E-F and 5). Tyr114 swings its side chain into the core of the protein, and makes a hydrogen bond with the side chain of Asn124, and interacts with the main chains of Leu102, Pro103 and Ala104, allowing the direct connection between the two ends of the RL2 loop. In addition, the main chain of Tyr114 also interacts with the side chain of Asn74 located in the RL1 loop of the same protomer. The replacement of Tyr114 by alanine displayed a drastic impact on the enzymatic activity, resulting in a reduced activation by FBP and a strong impairing effect in AMP-mediated inhibition.^41^ Tyr114 and its equivalent residue Phe117 in *Pt*AGPase^33^ lay in the proximity of the ATP adenine motif, suggesting that the observed effect is due to the impairment of ATP binding.^37^ The RL2 conformation is further stabilized by two hydrogen bonds between the main chain of Ala104 and the side chain of Asn112, and that of the side chain of Gln106 with the main chain of Glu111 (Figures 2C, 3C, and 5). This structural arrangement is reinforced by the interaction between the main chain of Trp113 and the side chain of Gln74, located in the RL1 loop. Moreover, the side chain of Gln105 interacts with the side chain of Gln76, located in the RL1’ loop of a neighbor subunit. Interestingly, the side chain of Arg107 makes a hydrogen bond with the main chain of Asn38, located in the SM motif. As a consequence, Arg29 swings its side chain out the ATP binding site and makes a strong electrostatic interaction with the side chain of Thr37. Similarly, the side chain of Trp113 leaves out the catalytic pocket and adopt a completely different conformation. The replacement of Trp113 and Gln74 by alanine leads to a remarkable reduction in the activity and abolished the activation by FBP and inhibition by AMP.^37,32^ Furthermore, beyond their impact in the enzyme activity, the single point mutation to alanine of RL2 residues 105-111 reduce AMP inhibition.^32^ It is worth noting that RL2 residues are mainly conserved among AGPases from different sources, suggesting a common inhibition mechanism (Figure 6).

### A conformational switch modulates ATP binding

Our cryoEM studies on *Ec*AGPase-FBP and *Ec*AGPase-AMP complexes support the occurence of two conformational states that operate in a concerted manner to allosterically modulate the enzymatic activity of the enzyme. A ‘locked’ state where the binding of AMP promotes the close interaction between the RL2 and RL2’ from different protomers of different dimers, markedly reducing their availability for ATP binding (Figure 5, Video S2: https://www.dropbox.com/s/6fu2cj8won6uu0i/vf_Video_S2.avi?dl=0); FBP displaces the negative regulator from the regulatory site and triggers the release of the RL2 and RL2’ loops adopting an ‘free’ state (Figures 4E-F). This conformational change makes the RL2 and RL2’ loops available for the interaction with ATP at the corresponding active sites. As a consequence, FBP breaks the symmetry of the particle, arguably from an *Ec*AGPase-AMP_D2_ inhibited state to an *Ec*AGPase-FBP_quasyC2_ active state, making the ATP site available. Interestingly, the C2 symmetric nature of the active enzyme can be assumed from early studies on *Ec*AGPase.^42^ This mechanism accounts for the fact that sensitivity to AMP inhibition is modulated by the concentration of the activator FBP.^19,34^

To determine whether the allosteric regulators modulate the recognition of ATP at the active site, we first determined the accessibility of the active site to substrates/products, in the presence/absence of the allosteric regulators by using a thermofluor assay (Figure 7A; please see Materials and Methods in Supplementary Information). In the low temperature range (4 to 45 °C), differences in fluorescence reflect the relative accessibility of the active and/or regulatory sites to the fluorophore SyPRO orange and therefore are attributable to the occupancy of the corresponding sites of *Ec*AGPase in the native state. A reduction of the fluorescence signal in the presence of ADP-glucose is assigned to the occupancy of the active site. Conversely, differences in the fluorescence signal in the presence of allosteric regulators indicate the occupancy of the allosteric cleft. As depicted in Figure 3E, the reduced fluorescence profile of *Ec*AGPase-AMP compared with the *Ec*AGPase-FBP complex strongly suggests greater accessibility of the active site to the fluorophore, and possibly the partial occupancy of the allosteric site by FBP. Moreover, in the high temperature denaturation range (45 to 95 °C), the shift in the melting temperature clearly support the stabilization of the enzyme mediated by AMP association.^16,30^ Interestingly, AMP binding prevented the loss of enzymatic activity by chemical modification of the active site.^43^ Finally, to determine the binding of ATP to the *Ec*AGPase-FBP and *Ec*AGPase-AMP complexes, we performed ITC experiments. The titration of the *Ec*AGPase-FBP with ATP discloses a very high affinity for the substrate, whereas, in contrast, the affinity towards ATP by the *Ec*AGPase-AMP complex is markedly reduced (Figure 7B; Figure S8; Table S1), supporting our proposed allosteric model for *Ec*AGPase (Figure 8).

### The allosteric network for signal transmission

To elucidate the pathways for the transmission of the positive and negative allosteric signals, we analyzed the residue interaction network (RIN) of (i) the FBP and AMP regulators, and (ii) the RL2 loop, towards the central core of the *Ec*AGPase homotetramer, respectively. As depicted in Figure 4, there are important differences in the RIN profiles observed in the cryoEM *Ec*AGPase-FBP and *Ec*AGPase-AMP complexes. The negative regulator AMP interacts with Thr79 (α5), in close contact with His78 (α5), which in turn associates with Gln105 located in the RL2 loop adopting a ‘locked’ conformation (Figure 5). In contrast, in the presence of the positive regulator FBP, the interaction between Gln105 and His78 is missing, modifying the communication with the RL2 loop which adopts an ‘free’ conformation (Figure 5). These differences also impact the RIN associated with the CL loop. Notably, the CL loop in the *Ec*AGPase-FBP complex encompasses a larger RIN (Figures S9-S10), compared to that observed in the *Ec*AGPase-AMP complex, including the coordination between Arg67 and its counterpart in the GT-A-like domain of the opposite protomer. The replacement of Arg67 by alanine caused changes in the allosteric properties of *Ec*AGPase.^44^ Furthermore, the CL interacts with a long loop that runs at the base of each protomer connecting the GTA-like domain and the LBH, here called the Inter-Domain Loops (IDL, residues 295-316). Interestingly, the CL interacts with the IDL from the same protomer and with both IDLs from the opposite dimer. The IDL anchors several regulatory motifs from the same and other protomers (SM, CL, IDL, LBH), changing its RIN in the FBP and AMP complexes. The IDL being at the core of the protein, therefore, represent a major element for allosteric signal transmission (Figures S9-S10).

## DISCUSSION

Bacterial glycogen and plant starch represent internal deposits of environmental carbon and energy surplus harvested by organisms.^1,5^ In the case of circumstantial scarcity, these storages are the source of energy permitting cell functionality. From an evolutionary perspective, the rise of glycogen metabolism results in a competitive advantage to these organisms. This functionality requires the control of the energy flow towards and from the glycogen storage. Thus, a signal-control system tightly decouples synthesis and degradation in time and space, adjusting the energy flux of the glycogen compartment to the central energy metabolism. In bacteria, AGPase controls glycogen biosynthesis, whereas, GP regulates glycogen degradation.^5^ The coordination of these pathways is often evident, as in the case of *E. coli*, where AMP mediates the inhibition of AGPase and the activation of GP. In addition, ADP-glucose acts as a competitive inhibitor of GP in *E. coli.* This archetypal example demonstrates that allosteric enzymes are one of the finest products of evolution, providing rapid mechanisms of regulation and signal coordination of the whole metabolism.

Upon the addition of AMP to *apo Ec*AGPase, the apparent melting temperature (*Tm*) value significantly increased, supporting the stabilization of the enzyme. In contrast, the addition of FBP to *apo Ec*AGPase, did not affect the *Tm*, even at a high concentration of the positive regulator. The addition of FBP to the *Ec*AGPase-AMP complex triggered a clear reduction in the *Tm* value, indicating that FBP compete with AMP and modify the structural arrangement of the *Ec*AGPase-AMP complex.^16^ Looking for differences in the secondary structure of *Ec*AGPase in the presence or absence of allosteric regulators in solution, we assessed the circular dichroism spectrum of the enzyme in the far-UV region, from 200 to 250 nm (Figure S11). Spectra were measured in a range of temperatures, from 10 to 40°C. Overall, the spectral curves are similar, indicating that there are no large changes in the secondary structure content triggered by the regulators. Nevertheless, a close inspection of the 205-220nm region shows slight differences supporting that *Ec*AGPase resembles (i) more the AMP-state at lower temperatures, and (ii) more the FBP-state at high temperature. Interestingly, enzyme kinetics measurements revealed that the addition of AMP to the *Ec*AGPase-FBP complex drastically reduced the enzymatic activity, compared to *Ec*AGPase in the absence of the positive allosteric regulator.^42^ Altogether, the biochemical and biophysical observations suggest that, in the absence of allosteric regulators, the structural arrangement of the active site of *Ec*AGPase is similar and/or visits more often the ‘locked’ conformation observed in the *Ec*AGPase-AMP complex. This view hints that the role of AMP is to stabilize a pre-existing low activity conformation already available in *apo Ec*AGPase. In turn, FBP displaces AMP inducing a change in the RIN that ends with the free activated ‘free’ conformation of the RL2 loops in the active site (Figure 5; Figure S10).

Kinetic properties of *Ec*AGPase single-point mutants have been previously reported.^32,37,38^ The harmonization of these results from different conditions and the interpretation of the kinetics parameters in the context of the structural data provided by X-ray crystallography and cryoEM represent a challenge. To reduce the complexity, we have performed a simplified meta-analysis of *Ec*AGPase mutants restricted to the replacement of selected residues by alanine (Table 2; please see Materials and Methods in Supplementary Information). In the absence of AMP and FBP, the kinetics parameters Vmax and S0.5 (ATP concentration yielding Vmax/2; Table 2) did not significantly change in *Ec*AGPase mutants of the active site. In contrast, the *Ec*AGPase variations Arg40Ala, Arg52Ala, and Arg386Ala, localized in the allosteric cleft of the enzyme, showed a markedly reduction in Vmax, suggesting that the positively charged pocket is required to maintain the functional structure of the SM (Figures 2-5). Moreover, Arg40Ala exhibits an increased S(ATP)0.5, supporting a possible impact in the neighboring catalytic residue Lys42 (Figure S1).

**Table 2:** Qualitative meta-analysis of the enzymatic properties of *Ec*AGPase single point mutants relative to wild-type.

|  | Residue location | Element | Mutant | basal (*apo*) |  | Activation (FBP) |  |  | Inhibition (AMP and FBP) | %Ident | Conserv score |
| --- | --- | --- | --- | --- | --- | --- | --- | --- | --- | --- | --- |
|  |  |  |  | Vmax ratio | S<sup>ATP</sup>(0.5) ratio | Vmax ratio | S<sup>ATP</sup>(0.5) ratio | A(0.5) ratio | I(0.5) ratio |  |  |
| N-term domain (GTA-like) | Active site | RL2(a) | P103A | ~ | ↓ | ↓ | ↑ | ~ | ~ | 75 | 7 |
|  |  |  | Q105A | ~ | ~ | ~ | ↑ | ↑ | ↑ | 75 | 6 |
|  |  |  | **Q106A** | ~ | ~ | ↓ | ↑ | ↑ | ~ | **100** | **11** |
|  |  |  | R107A | ~ | ~ | ↓ | ↑ | ~ | ~ | 58 | 6 |
|  |  |  | M108A | ~ | ~ | ~ | ~ | ~ | ~ | 50 | 5 |
|  |  |  | K109A | ~ | ~ | ~ | ↑ | ~ | ~ | 33 | 2 |
|  |  |  | E111A | ~ | ~ | ~ | ~ | ~ | ~ | 41 | 2 |
|  |  |  | N112A | ~ | ~ | ~ | ↑ | ~ | ~ | 33 | 5 |
|  |  |  | **W113A**<br>* | ~ | ~ | ↓ | ↑ | ~ | ~ | **100** | **11** |
|  |  |  | **Y114A** | ~ | ~ | ↓ | ↑ | ~ | ~ | **66** | **9** |
|  |  |  | R115A | ~ | ~ | ↓ | ↑ | ↑ | ~ | 41 | 4 |
|  |  | RL1(b) | **Q74A** | n/a | ~ | n/a | ↑ | n/a | n/a | **100** | **11** |
|  | Regulator site (c) | SM | **R40A** | ↓ | ↑ | ↓ | ↑ | n/a | n/a | **91** | **8** |
|  |  |  | **R52A** | ↓ | ~ | ↓ | ↑ | n/a | n/a | **100** | **11** |
|  |  | α-7 | R130A | ↑ | ~ | ~ | ~ | ↑ | n/a | 33 | 1 |
| C-term domain (LBH) |  | LBH | R353A | ~ | ~ | ~ | ~ | ~ | n/a | 58 | 2 |
|  |  |  | R386A | ↓ | ~ | ↓ | ↑ | ↑ | n/a | 50 | 5 |
|  |  | C-term α-helix | R419A | ~ | ~ | ~ | ~ | ~ | n/a | 33 | 2 |
|  |  |  | R423A | ~ | ~ | ~ | ~ | ↑ | n/a | 33 | 2 |

a. Reference 32
b. Reference 37
c. Reference 38
\*also reported in b

In the FBP-activated state, mutants localized both in the active and regulatory sites of *Ec*AGPase showed altered kinetics profiles (Table 2). In particular, mutants Pro103Ala, Gln106Ala, Arg107Ala, Trp113Ala, Tyr114Ala and Arg115Ala, located in the RL2 loop, are primarily characterized by a lack of response to FBP (Figure 5; Table 2).^32^ Specifically, Pro103Ala displayed a reduced Vmax and reduced affinity for ATP. Pro103 is a strictly conserved residue that appears to play a fundamental role in the structural arrangement of the RL2 loop. Gln106Ala also displayed a reduced Vmax and affinity for ATP, and required a higher concentration of FBP to activate the enzyme. In the *Ec*AGPase-FBP_D2_ complex, the side chain of Gln106 makes strong interactions with the main chain of Arg115, located in the RL2 loop, and Gln74, located in the RL1 loop (Table 2). In contrast, in the *Ec*AGPase-AMP_D2_ complex, the side chain of Gln106 displays a completely different arrangement, making a hydrogen bond with the main chain of Gln112. The Arg107Ala mutant displays a reduced Vmax and affinity for ATP. The side chain of Arg107 makes an electrostatic interaction with the side chain of Gln123, stabilizing the ‘free’ conformation in *Ec*AGPase-FBP_D2_ complex. Arg107 is located in *α*7, a key secondary structure that mediates the signal transmission between the regulatory site and the active site through the RL2 loop. The Trp113Ala and Tyr114Ala mutants also showed a reduced Vmax and affinity for ATP. Although both residues have been suggested to participate in the binding of ATP, their side chains are not observed in the cryoEM structure of the *Ec*AGPase-FBP complex, limiting the analysis. Arg115 showed a reduced Vmax and affinity for ATP, and required a higher concentration of FBP to activate *Ec*AGPase. The side chain of Arg115 makes an electrostatic interaction with the side chain of His238, located in *α*7, contributing to the formation of the ‘free’ conformation of the RL2 loop in the *Ec*AGPase-FBP_D2_ complex. Finally, at the regulatory cleft, mutations Arg40Ala, Arg52Ala, and Arg386Ala displayed a reduced Vmax and S0.5 values, possibly by impairing the arrangement of the activator phosphate group in the corresponding pocket. Interestingly, although Arg130Ala and Arg423Ala display similar Vmax and S0.5 values; the mutants require higher concentrations of FBP to achieve half-full activation levels, an indication of their role in FBP recognition and the activation mechanism.

In the case the FBP-activated state was inhibited by AMP, all reported mutants located in the active site, including Pro103Ala, Gln106Ala, Arg107Ala, Met108Ala, Lys109Ala, Glu111Ala, Asn112Ala, Trp113Ala, Tyr114Ala, and Arg115Ala, did not display a substantial change in the I0.5, the AMP concentration required for half inhibition of the enzymatic activity of *Ec*AGPase, possibly due to the preservation of the AMP site architecture. The Gln105Ala represents an exception, requiring a higher concentration of AMP for inhibition. In the AMP-inhibited state, Gln105 interacts with the side chain of Gln76, located in the RL1 loop; and the main chain of His78, located in the top of *α*5, where Thr79 participate in the interaction with the *α* phosphate group of AMP. Interestingly, mutants of the RL2 loop which exhibited the least sensitivity to FBP were also the least sensitive to AMP inhibition. *Ec*AGPase was activated approximately 25-fold by FBP, compared to 1.5 fold for the mutants Pro103Ala, Trp113Ala, and Tyr114Ala. *Ec*AGPase conserved just ca. 3% of its enzymatic activity with saturating AMP, whereas mutants Pro103Ala, Trp113Ala, and Tyr114Ala retained ca. 50–70%. Altogether, the structural and enzymatic data support the notion that these residues, located at both ends of the RL2 loop, play a major mechanistic role to facilitate the ‘free-to-locked’ transitions, critical for the regulation of the enzyme (Figure 5). We previously reported that mutants Lys39Ala, Arg40Ala, His46Ala, Arg52Ala, Arg386Ala, Arg419Ala and Arg423Ala, located in the regulatory site, failed to achieve the activation values mediated by FBP, compared to that obtained for and severely compromised the inhibition by AMP. Strikingly, we found that the Arg130Ala mutant deregulated AMP-mediated inhibition of the enzymatic activity, inducing the overproduction of glycogen *in vivo*. The interaction of Arg130 (*α*7) with the AMP nucleobase stabilizes the *α*7 helix in such a way facilitating the arrangement of a cavity for the accommodation of Tyr114 in the locked conformation.

The enzyme cooperativity mechanism is disclosed by the *Ec*AGPase-FBP_D2_ and *Ec*AGPase-AMP_D2_ reconstructions, revealing clear pathways of communication between the active sites in the homotetrameric architecture. The signal transmission is mainly mediated by the interactions between the GT-A like domains (Figure 8; Figure S6). It occurs essentially in two manners: (i) side-by-side, where one GTA-like domain of a protomer interacts with a GTA-like domain of another protomer from a different dimer (Figure 8); and (ii) top-to-bottom, where the interactions between GTA-like domains occur across the tetramerization interphase (Figure S6). It has been extensively reported that FBP activation increases the positive cooperative effect displayed by *Ec*AGPase. Our study clearly shows that FBP interaction with *Ec*AGPase mediates the release of the RL2-to-RL2 lock, which means an immediate availability of two active sites in the active conformation. This event explains the drastic increment in the activity of the enzyme. Moreover, comparative RIN analysis between the AMP and FBP states reveal that the release of the “locked” conformation is accompanied by the reduction of the RL2 interactions. Simultaneously, there is an increment of the CL RIN, which is anchored to the *α*5, a central element in the architecture of AGPases. The *α*5 repositioning appears to contribute to the release of the RL1-RL2 interactions in the “locked” conformation. Finally, the changes in the CL conformation is disseminated across the tetramer by interactions with the IDL, which associates with several regulatory structural elements from different protomers.

Several residues involved in catalysis are conserved in the active site across all AGPase homologs, including (i) Arg32 involved in the anchoring of the γ phosphate group of ATP, and (ii) Lys42 and Lys195, involved in the interaction, respectively, with ATP and G1P, which polarize the phosphates groups to facilitate the nucleophilic reaction (G-rich loop; Figure S1). Moreover, several residues implicated in substrate recognition are also preserved (Figure 6). In contrast, most residues at the regulatory cleft implicated in the binding of FBP and AMP regulators in *Ec*AGPase are not highly conserved, with the exception of Arg40 and Arg52 (Figure 6). This observation agrees with the fact that AGPases from different sources use different allosteric regulators, providing a specific relationship to specific metabolic routes of relevance to the organism or tissue. The conservation of Arg40 and Arg52 can be explained because both residues participate in the interaction with the *α* phosphate group of ATP or a phosphate moiety of FBP, preserving the positively charged pocket shaped by the SM. This pocket might participate in the binding of other phosphorylated metabolites known to regulate other AGPases, as 3-phosphoglycerate (3PGA) in plants or fructose-6-phosphate (F6P) in *Agrobacterium tumefaciens*.^17,31^ The long RL2 loop appears to be an evolutionary feature of AGPases, not observed in other non-regulated Nucleotide-di-phosphate pyrophosphorylases (NDPases; Figure S12). The structural characteristics, the close location to the ATP substrate, and the conservation of the RL2 loop strongly suggest this structural element as a major driving force in AGPase evolution (Figure 6, Table 2). Altogether, these facts imply the evolution of the allosteric cleft is partially disentangled from the evolution of the active site, facilitating the divergence of regulatory cleft residues to acquire different enzymatic regulatory characteristics.

Considered the ‘second secret of life’,^45^ allosterism is a central biological phenomenon that provides coherence and harmony to the metabolism. An allosteric enzyme is a specialized product of evolutionary engineering enabling the control of catalysis by ligands that are chemically foreign to the chemical reaction; therefore, it is the base of metabolic coordination. In words of Monod, “the very gratuitousness of these systems, giving molecular evolution a practically limitless field for exploration and experiment, enabled it to elaborate the huge network of cybernetic interconnections which makes each organism an autonomous functional unit, whose performances appear to transcend the laws of chemistry”.^13^

Different models have been proposed to contextualize the current understanding of the allosteric phenomena. The original Monod-Wyman-Changeux (MWC) concerted model presents a perspective of multimeric enzymes where all protomers are constrained to a single conformational state at a given moment.^46^ Two global conformational states exist in equilibrium, the so-called T (tense) and R (relaxed) states. The conformational transition between the two states occurs simultaneously, in a concerted manner, in all protomers. The ligands have different affinity for the two states, and the equilibrium is shifted thanks to the binding of an allosteric effector or substrate to a protomer. Since only one conformer can exist in a given enzyme molecule at the time, this model implies infinite coupling between protomers suppressing conformational intermediates.^47^ The R-T shifting mediated by ligand binding represents the view of a conformational selection of the conformers. According to the MWC model view, the *Ec*AGPase-AMP complex represents a T inhibited state since the ‘locked’ mechanism requires all protomers to acquire the same conformation (Figures 3, 4 and 5; Figure S6).^16,30^ This approximation appears feasible since the inhibitor is active at very reduced concentrations, pointing to the high affinity of AMP, leading to an almost instant full occupancy of the allosteric cleft according to the reported intracellular AMP concentration, even if low amounts of FBP are present.^48-51^

In the Koshland, Némethy and Filmer (KNF) sequential model, the protomers are not constrained to a global concerted conformational change of all protomers to the same conformation. Therefore, the KNF model contemplates the enzyme subunits to exist in different conformations at any given time.^52^ The binding of a substrate or effector to a protomer influence in a different degree the conformation of other protomers and their affinity towards the ligands. The KNF model implies that ligands bind via induced fit phenomena, triggering conformational changes. In the case of AMP displacement by FBP, we observe that the activated form of *Ec*AGPase shows protomers in different conformations, which concurs with this view. Moreover, our observations showing that FBP can have differences in the binding mode and differences in conformations are in agreement with the induced fit view.

Although these classical models of allosterism address some of the observations, the different conformations observed locally points towards other paradigms of allosterism. The *Ec*AGPase model of regulation can be better reflected by the Ensemble Model (EM) in which the allosteric system is described as a population of conformational states statistically distributed according to their energy levels.^53^ In that sense, single-particle cryoEM reflects the view of the consensus of the particles. Other alternative paradigms present the allosteric phenomena without large motions of the enzyme but instead with changes in the dynamic and minimum global conformational changes. According to this view, the binding of the allosteric effector transfers the allosteric signals to the active site in the form of entropy-driven dynamics.^54^ Importantly, this model has been proposed for kinases having similar G-rich loops to AGPases.^55^ Specifically, the transmission of the signal can be defined by two mechanisms: (i) a ‘domino model’ propagating the signal through sequential local structural changes from the allosteric site via a single pathway to the active site and (ii) a ‘violin model’ analogy mechanism, where the enzyme vibrational states, as the body of the violin, can change because of the binding of the effector, transferring the signal throughout the whole structure with no specific pathway.^55^

Our work highlights how the advances of cryoEM can extraordinarily power the study of the cooperative and allosteric phenomena of enzymes. We observe novel biologically relevant conformations of *Ec*AGPase that could not be visualized by other structural biology techniques – e.g. X-ray crystallography due to crystal constraints and crystallization artifacts. The single-particle reconstruction approach permits to build a model that represents the conformational consensus of the isolated enzyme. Strikingly, since the majority of allosteric and cooperative enzymes are multimeric proteins, the study of all allowed symmetries of the system reveal different views of the phenomena. In our case, although different symmetrized reconstructions result in a similar resolution, they allow different observations. In the activated state of *Ec*AGPase, the high symmetry D2 allows to model a consensus for the RL2 loop in the ‘free’ conformation, meanwhile, side chains conformation and the FBP occupancy of the allosteric cleft can be explored in the lower symmetries. On the other hand, all the reconstructions from the *Ec*AGPase-AMP complex disclose the same conformation, indicating that the single-particle reconstruction represents a more homogeneous ensemble of particles. Many more allosteric enzymatic systems await for being explored or re-visited by cryoEM to reveal their mechanisms.

## AUTHOR CONTRIBUTIONS

J.O.C. and M.E.G. conceived the project. A.M., N.C., and C.D. expressed and purified the protein. J.O.C and D.G.C. performed cryoEM grid optimization and cryoEM microscopy screening. J.O.C. and D.G.C. collected EM data. J.O.C. performed EM processing, built and refined the atomic models. J.O.C., D.A.J, N.C. and C.D. performed biophysical studies. J.O.C., C.D., N.C. and M.E.G. prepared figures and tables. J.O.C and M.E.G wrote the manuscript. All the authors critically discussed the interpretation of the data and the final manuscript.

## ACKNOWLEDGMENTS

This project was supported by grants from (i) the MINECO/FEDER EU contracts BFU2016-77427-C2-2-R, BFU2017-92223-EXP and Severo Ochoa Excellence Accreditation SEV-2016-0644, and (ii) the Basque Government contract KK-2019/00076 (to M.E.G.). We acknowledge Diamond Light Source for access and support of the cryo-EM facilities at the UK’s national Electron Bio-imaging Centre (eBIC) under proposals EM6916 and EM17171, funded by the Wellcome Trust, MRC and BBRSC. We thank for valuable assistance in cryoEM data collection to the eBIC Diamond staff, especially to Alistair Siebert, Corey Hecksel and Kyle Dent.

## SUPPORTING INFORMATION

### SUPPLEMENTARY TABLES

**Table S1.** Fitting parameters for ITC data for four sites ATP sequential binding model.

| Binding site | *EcAGPase*-FBP vs ATP |  | *EcAGPase*-AMP vs ATP |  |
| --- | --- | --- | --- | --- |
|  | K<sub>D</sub> (M) | ΔH (kcal/mol) | K<sub>D</sub> (M) | ΔH (kcal/mol) |
| 1 | $2.30 \times 10^{-9} \pm 6.48 \times 10^{-10}$ | $-7.93 \pm 1.60$ | $1.53 \times 10^{-4} \pm 2.22 \times 10^{-3}$ | $-28.5 \pm 676$ |
| 2 | $4.20 \times 10^{-7} \pm 4.36 \times 10^{-6}$ | $-2.85 \pm 25.7$ | $1.24 \times 10^{-3} \pm 1.94 \times 10^{-1}$ | $1.62 \pm 1130$ |
| 3 | $3.30 \times 10^{-10} \pm 3.42 \times 10^{-9}$ | $-0.86 \pm 24.2$ | $84.7 \times 10^{-9} \pm 21.6 \times 10^{-1}$ | $21.6 \pm 1360$ |
| 4 | $3.37 \times 10^{-4} \pm 1.15 \times 10^{-6}$ | $-29.9 \pm 4.32$ | $1.03 \times 10^{-3} \pm 1.51 \times 10^{-1}$ | $-40.4 \pm 7260$ |

### SUPPLEMENTARY FIGURES

**Figure S1.**
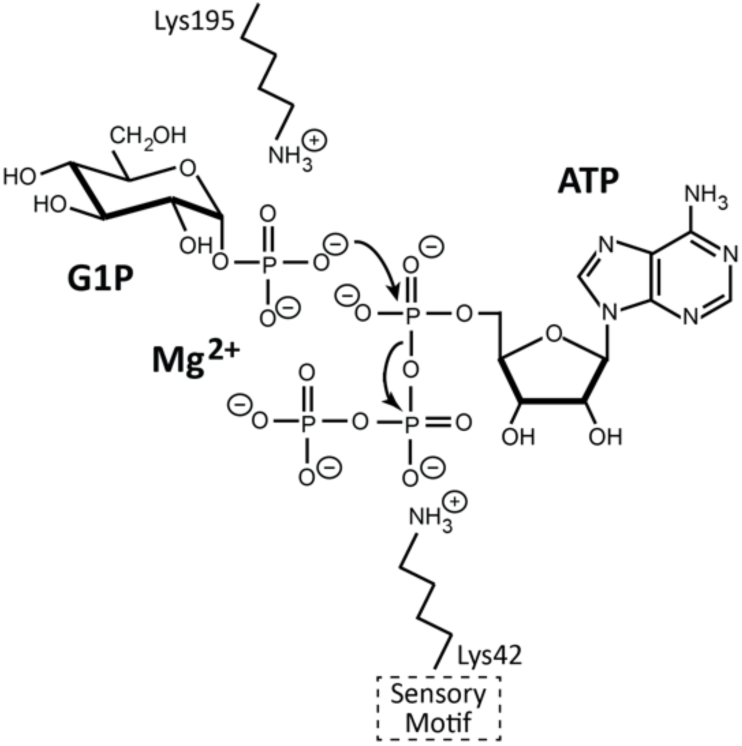
The catalytic mechanism of *Ec*AGPase. The α-phosphate from G1P must face the α-phosphate from ATP in the right geometry for the nucleophilic attack to take place. The positioning and the activation of the intervening phosphate groups relies on: (i) the interaction of the PO_4_^3-^ groups with the Mg^2+^ bound to the putative aspartate dyad, (ii) the anchoring of the ATP to the G-rich loop, with the γPO_4_^3-^ interacting with a highly conserved arginine, Arg32 in *Ec*AGPase, (iii) the interaction with two highly conserved catalytic lysines, Lys42 for ATP and Lys195 for G1P in *Ec*AGPase, which polarize the phosphates groups to facilitate the nucleophilic reaction. Finally, structural evidences in related enzymes support the ordered release of PPi first and ADPG second after the catalysis. The PPi molecule appears more solvent exposed and ready to leave the active site while the sugar phosphate is buried in the active site.

**Figure S2.**
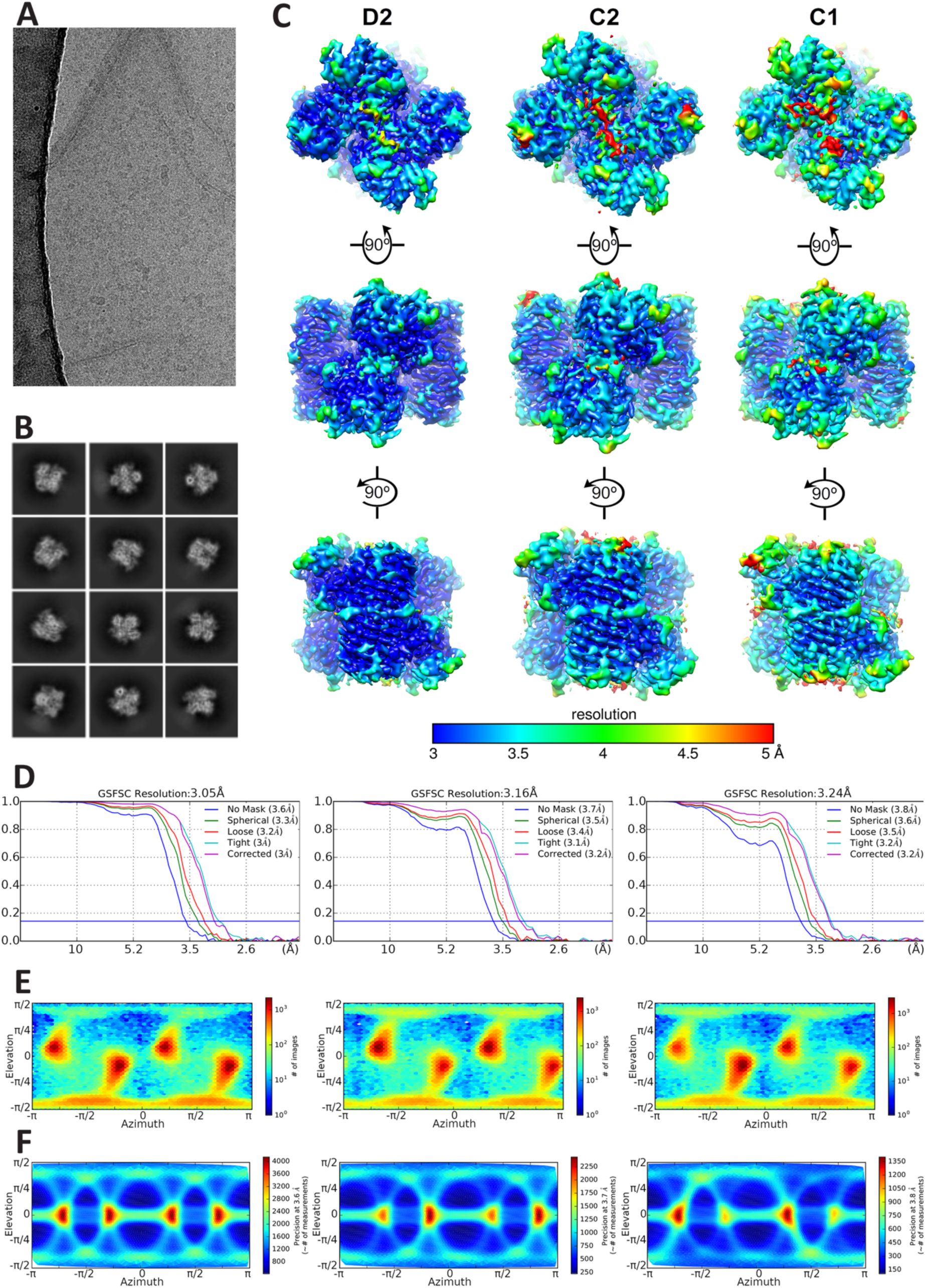
The *Ec*AGPase-FBP complex. (**A**) Micrograph detail. (**B**) Representative 2D classes. (**C**) Overall view of the single-particle reconstructions of *Ec*AGPase-FBP complex in D2, C2 and C1 symmetries, colored according to local resolution. (**D**) From left to right, the FSC plots of the *Ec*AGPase-FBP for the D2, C2 and C1 reconstructions. (**E**) From left to right, direction distribution of *Ec*AGPase-FBP particle images in the D2, C2 and C1 reconstructions. (**F**) From left to right, posterior precision plots for the *Ec*AGPase-FBP particle images in the D2, C2 and C1 reconstructions.

**Figure S3.**
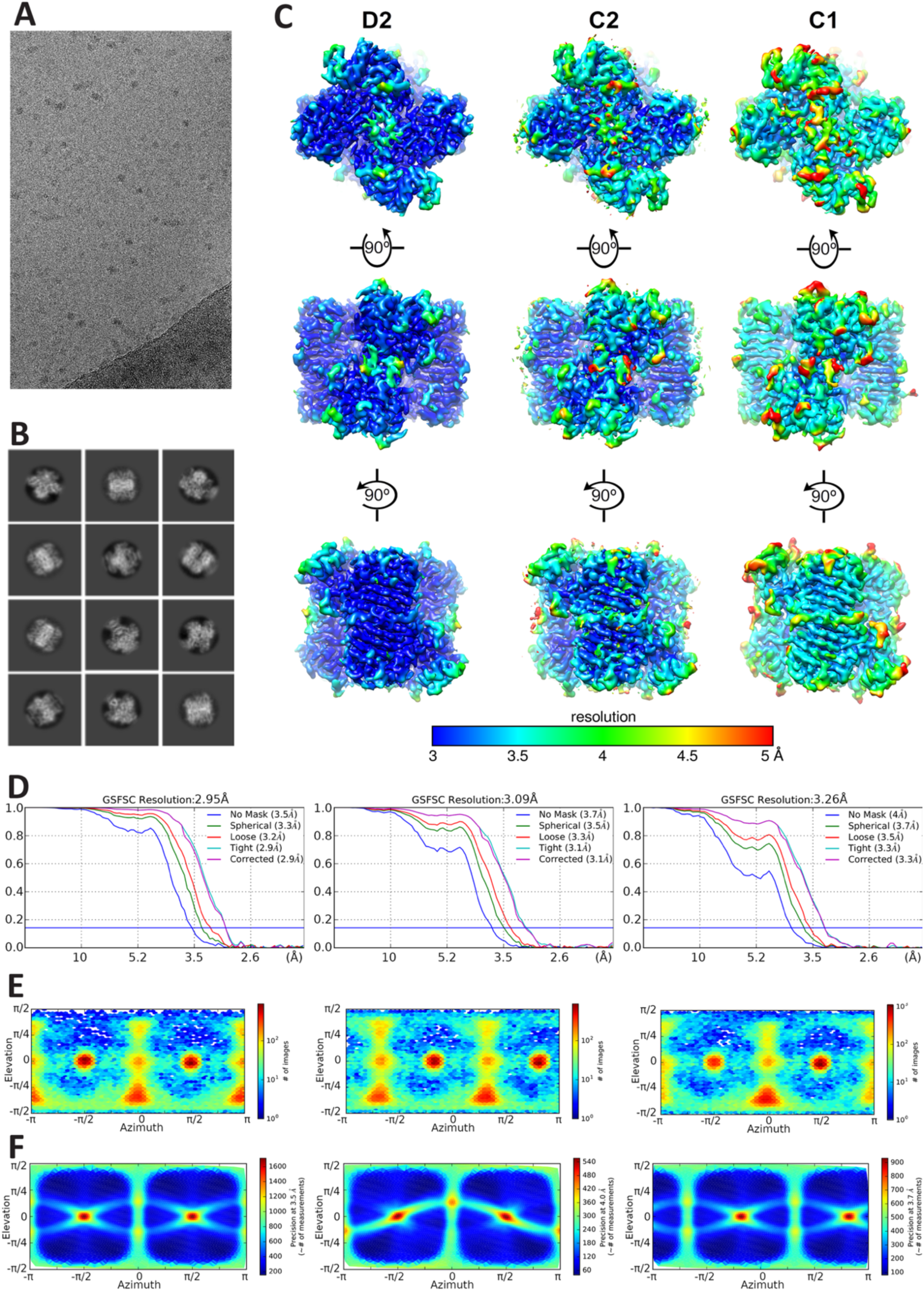
The *Ec*AGPase-AMP complex. (**A**) Micrograph detail. (**B**) Representative 2D classes. (**C**) Overall view of the single-particle reconstructions of *Ec*AGPase-AMP complex in D2, C2 and C1 symmetries, colored according to local resolution. (**D**) From left to right, the FSC plots of the *Ec*AGPase-AMP for the D2, C2 and C1 reconstructions. (**E**) From left to right, direction distribution of *Ec*AGPase-AMP particle images in the D2, C2 and C1 reconstructions. (**F**) From left to right, posterior precision plots for the *Ec*AGPase-FBP particle images in the D2, C2 and C1 reconstructions.

**Figure S4.**
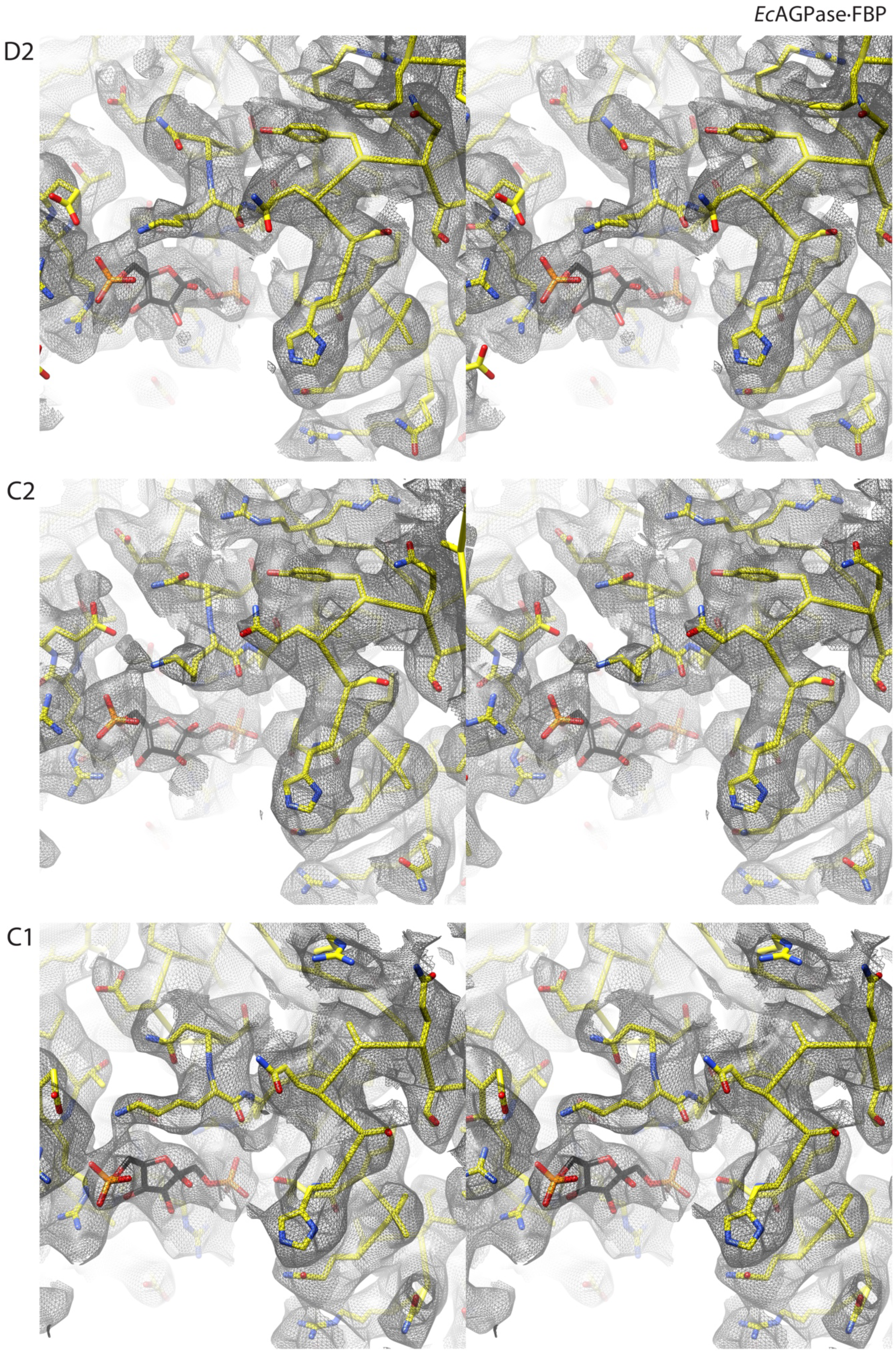
Local stereo views of the *Ec*AGPase-FBP model and maps. From top to bottom, a closed view of FBP ligand and the surrounding region of the D2, C2 and C1 map reconstructions and their corresponding models represented as yellow sticks.

**Figure S5.**
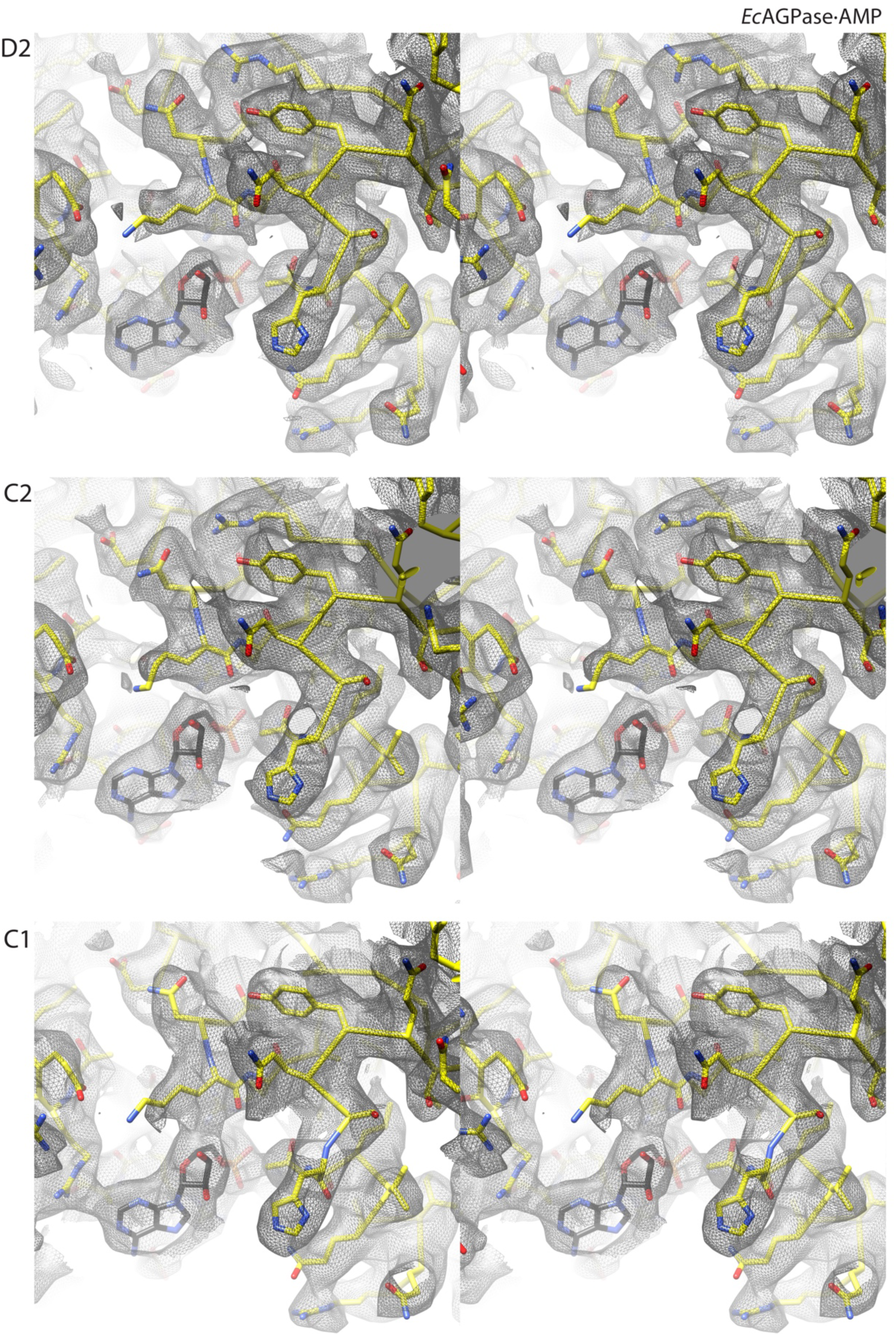
Local stereo views (crossed-eyes) of the *Ec*AGPase-AMP model and maps. From top to bottom, a closed view of AMP ligand and the surrounding region of the D2, C2 and C1 map reconstructions and their corresponding models represented as yellow sticks.

**Figure S6.**
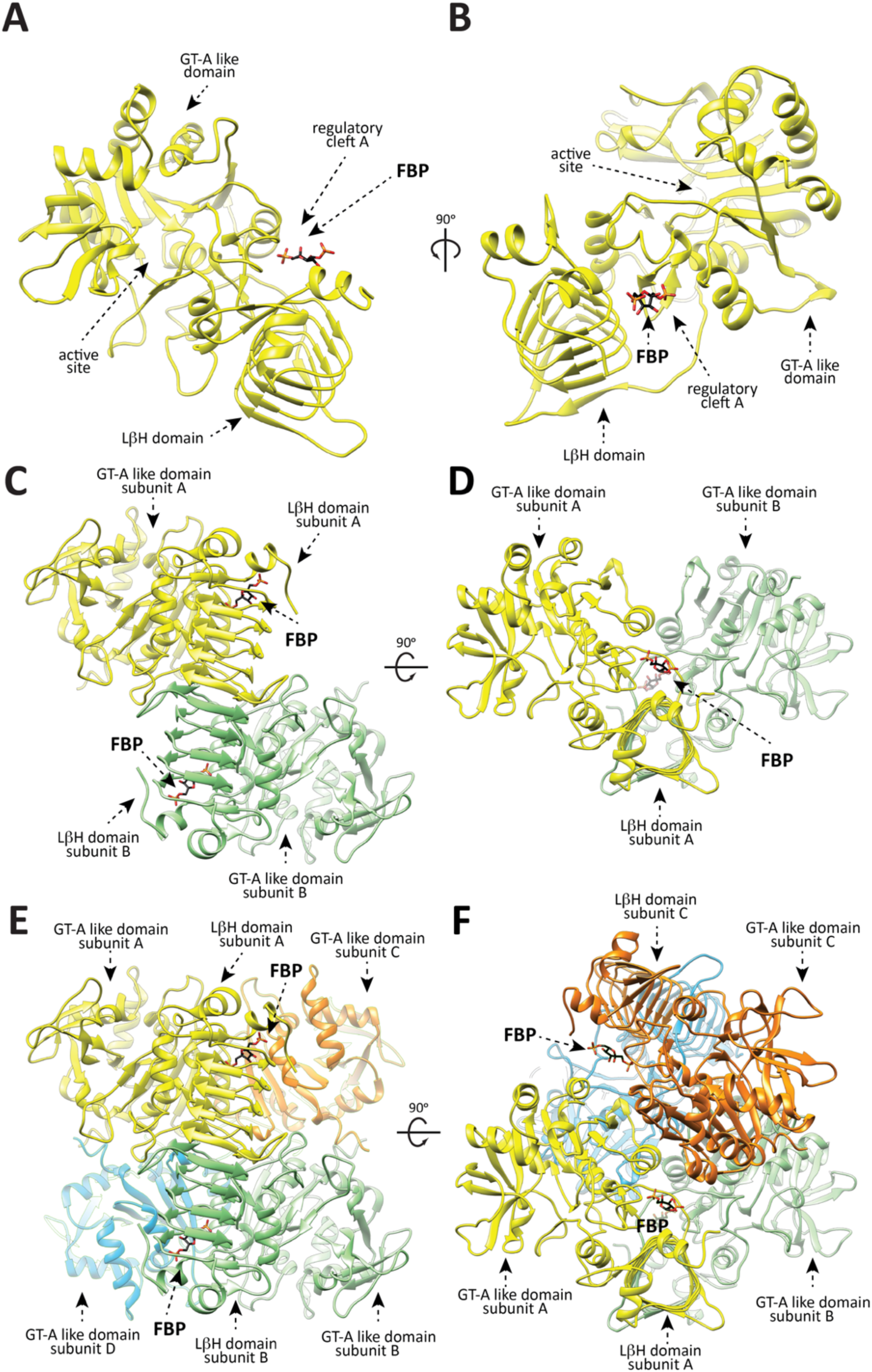
Overall structure of *Ec*AGPase as visualized by CryoEM. (**A**) Two views of an *Ec*AGPase protomer as visualized in the cryoEM *Ec*AGPase-FBP complex showing the GT-A-like domain and the LβH. (**B**) Two views of the *Ec*AGPase dimer. The protomers A and B are shown in yellow and green, respectively. (**C**) Two views of the *Ec*AGPase homotetramer. The protomers A, B, C and D are shown in yellow, green, orange and blue, respectively. The location of the active and regulatory sites is indicated.

**Figure S7.**
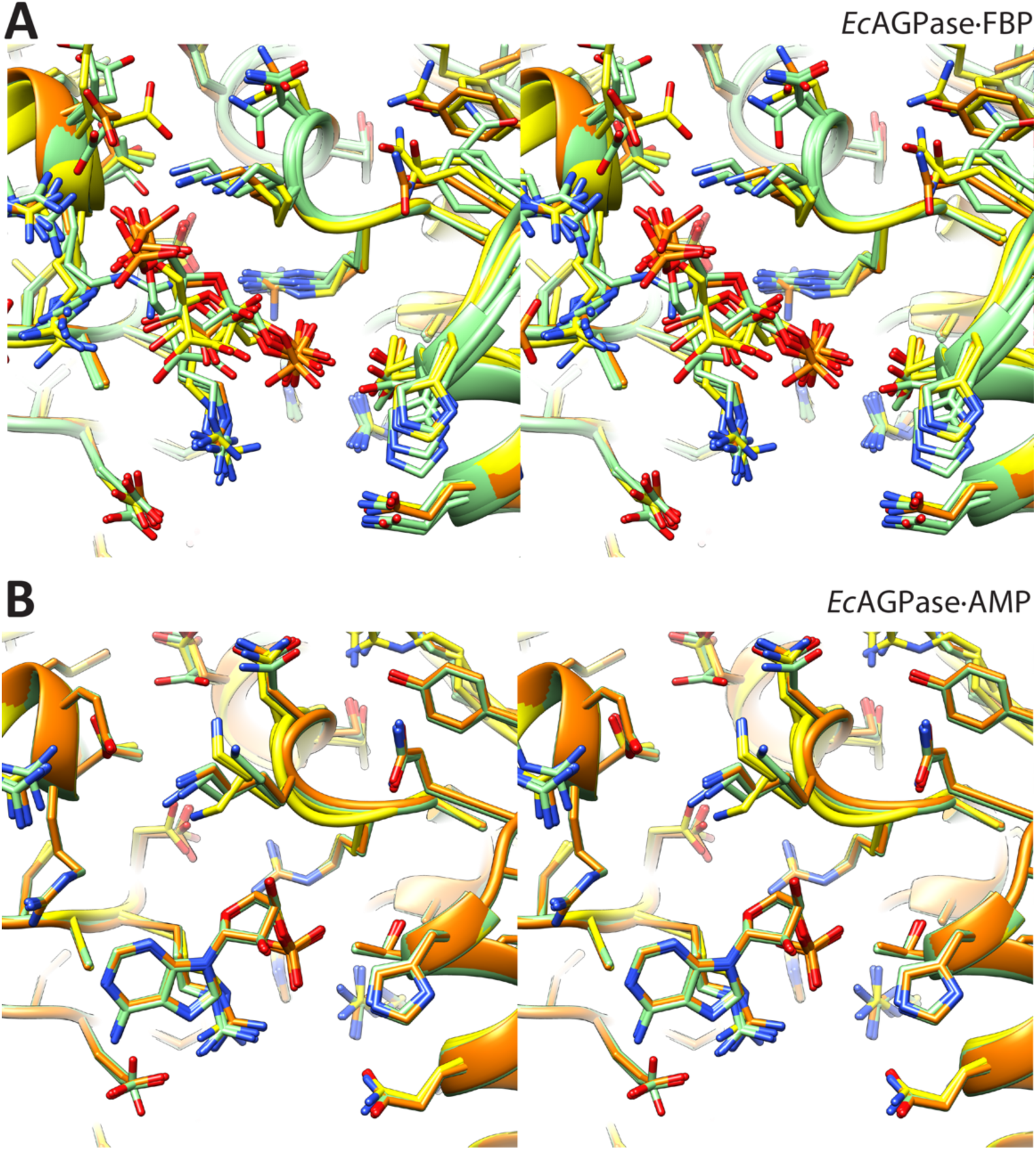
Local stereo views (crossed-eyes) of the allosteric cleft of *Ec*AGPase-FBP and *Ec*AGPase-AMP complexes. (**A**) Superposition of the D2 (orange), C2 (yellow), and C1 (green) models of the *Ec*AGPase-FBP complex showing the FBP ligand and the surrounding region. (**B**) Superposition of the D2 (orange), C2 (yellow), and C1 (green) models of the *Ec*AGPase-AMP complex showing the AMP ligand and the surrounding region.

**Figure S8.**
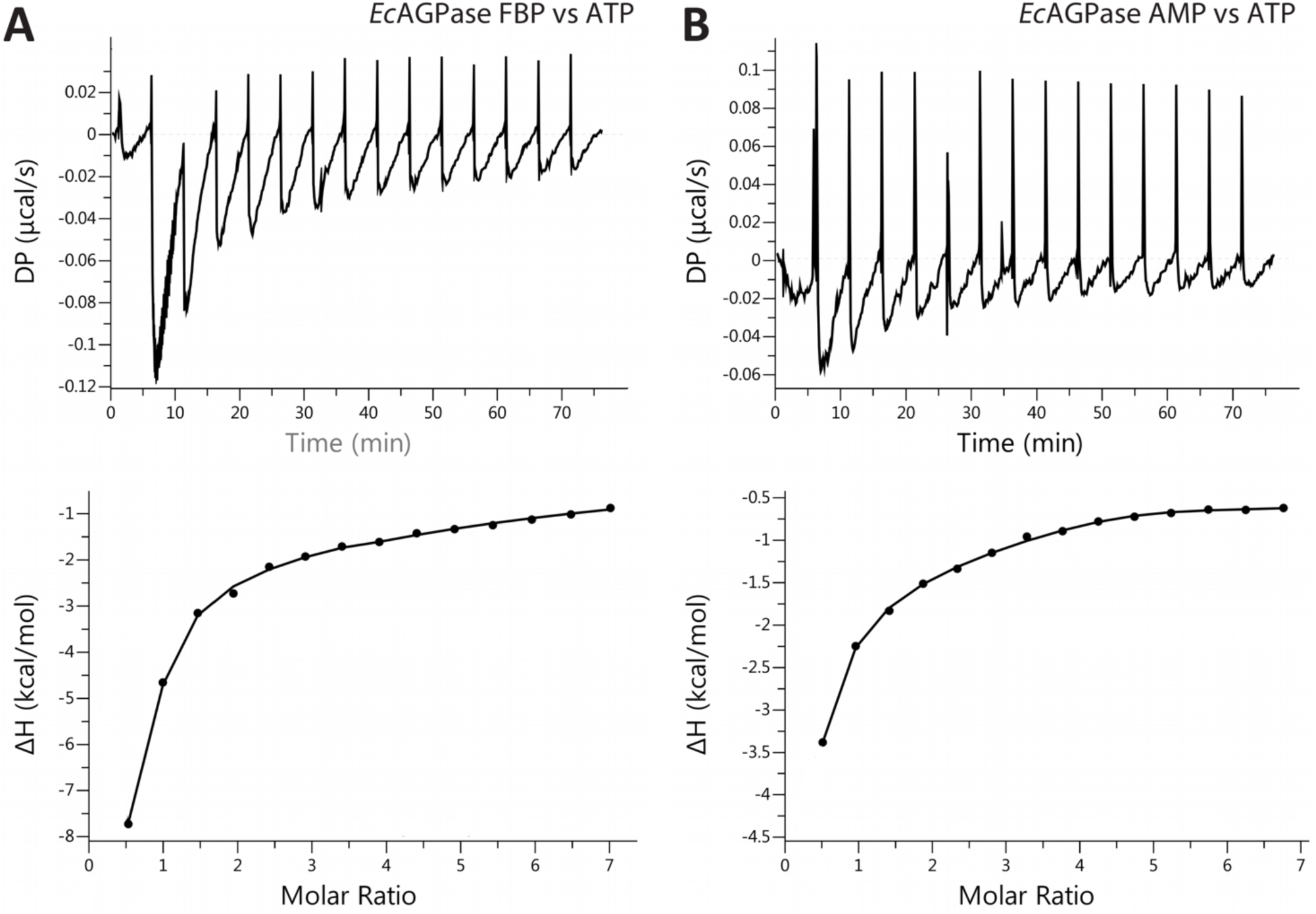
ITC measurements of *Ec*AGPase-FBP and *Ec*AGPase-AMP complexes with ATP. (**A**) The *upper panel* shows the raw data of the titration of the *Ec*AGPase-FBP complex with ATP. The lower panel shows the integrated heat of injections of the above titrations normalized per mol of ATP versus the molar ratio of ATP/*Ec*AGPase fitted to a sequential binding sites model. (**B**) The *upper panel* shows the raw data of the titration of the *Ec*AGPase-AMP complex with ATP. The lower panel shows the integrated heat of injections of the above titrations normalized per mol of ATP versus the molar ratio of ATP/*Ec*AGPase fitted to a sequential binding sites model.

**Figure S9.**
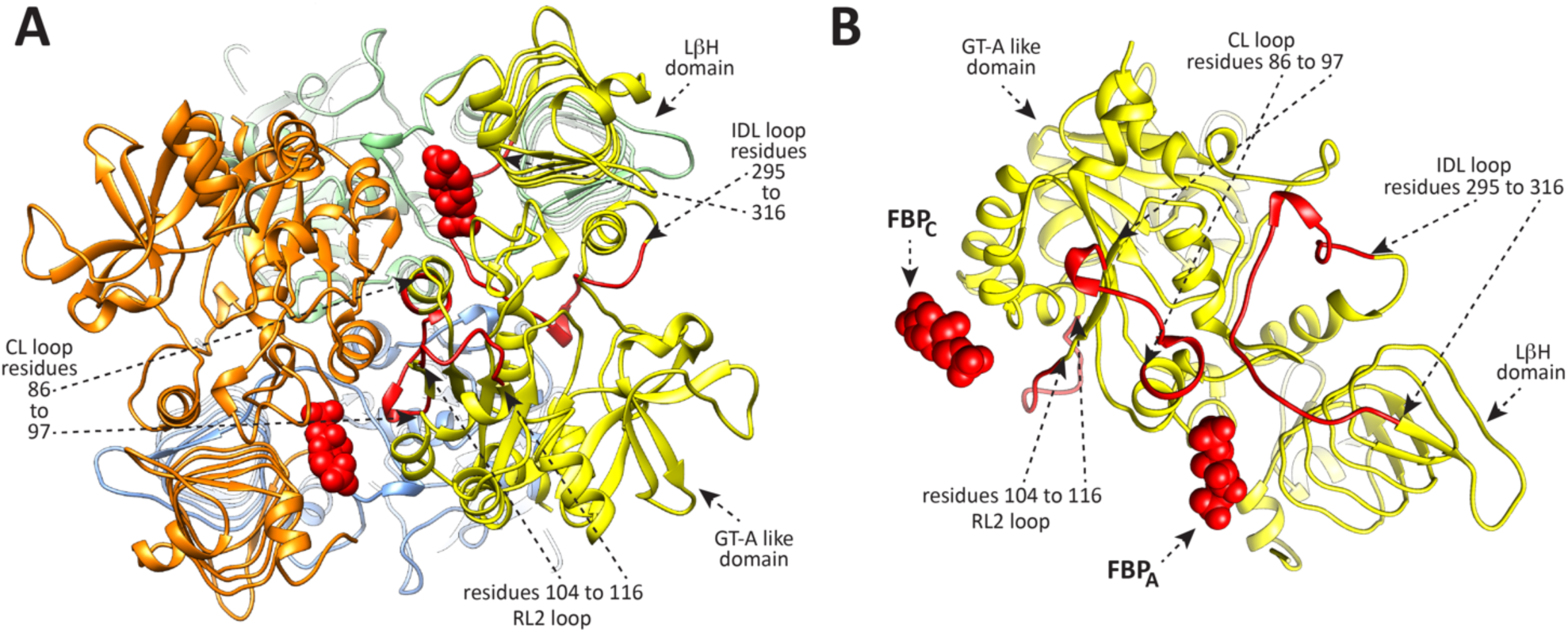
Allosteric residue interaction network (RIN) analyzed. (**A-B**) Two views of the *Ec*AGPase-FBP_D2_ model showing the RINs analyzed.

**Figure S10.**
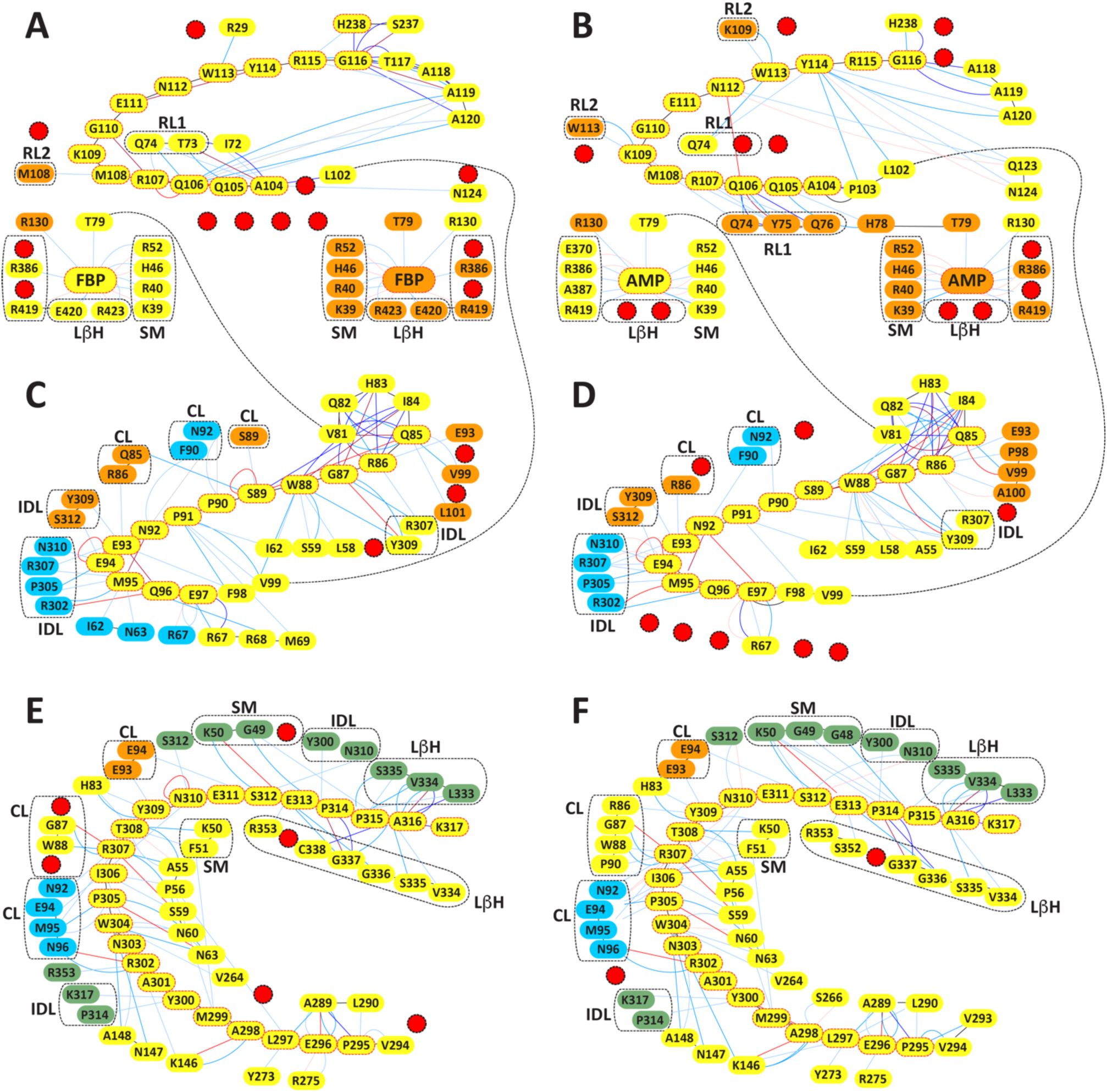
RIN differences between *Ec*AGPase-FBP and *Ec*AGPase-AMP complexes. (**A-B**). The RIN between the RL2 (104-116) and the corresponding allosteric regulator observed in the *Ec*AGPase-FBP_D2_ and *Ec*AGPase-AMP_D2_ complexes, respectively, are depicted as boxes highlighted in red. (**C-D**). The RIN between the CL (85-97) and the corresponding allosteric regulator observed in the *Ec*AGPase-FBP_D2_ and *Ec*AGPase-AMP_D2_ complexes, respectively, are depicted as boxes highlighted in red (**E-F**). The RIN between the IDL (104-116) and the corresponding allosteric regulator observed in the *Ec*AGPase-FBP_D2_ and *Ec*AGPase-AMP_D2_ complexes, respectively, are depicted as boxes highlighted in red. For panels (**A**) to (**F**), the interacting residues in other protomers are colored according to the panel (**A**) scheme described in Figure S9. The interactions are shown as straight or curved lines connecting residues. Interaction lines are colored according to different types of interactions as interatomic contacts (shades of blue), hydrogen bond (shades red), main chain bonds (black) and distant main chain connections (black dashed lines). Groups of residues from different elements are enclosed in black boxes. Red circles indicate missing residues in the interaction network between the *Ec*AGPase-FBP_D2_ and the *Ec*AGPase-AMP_D2_ models.

**Figure S11.**
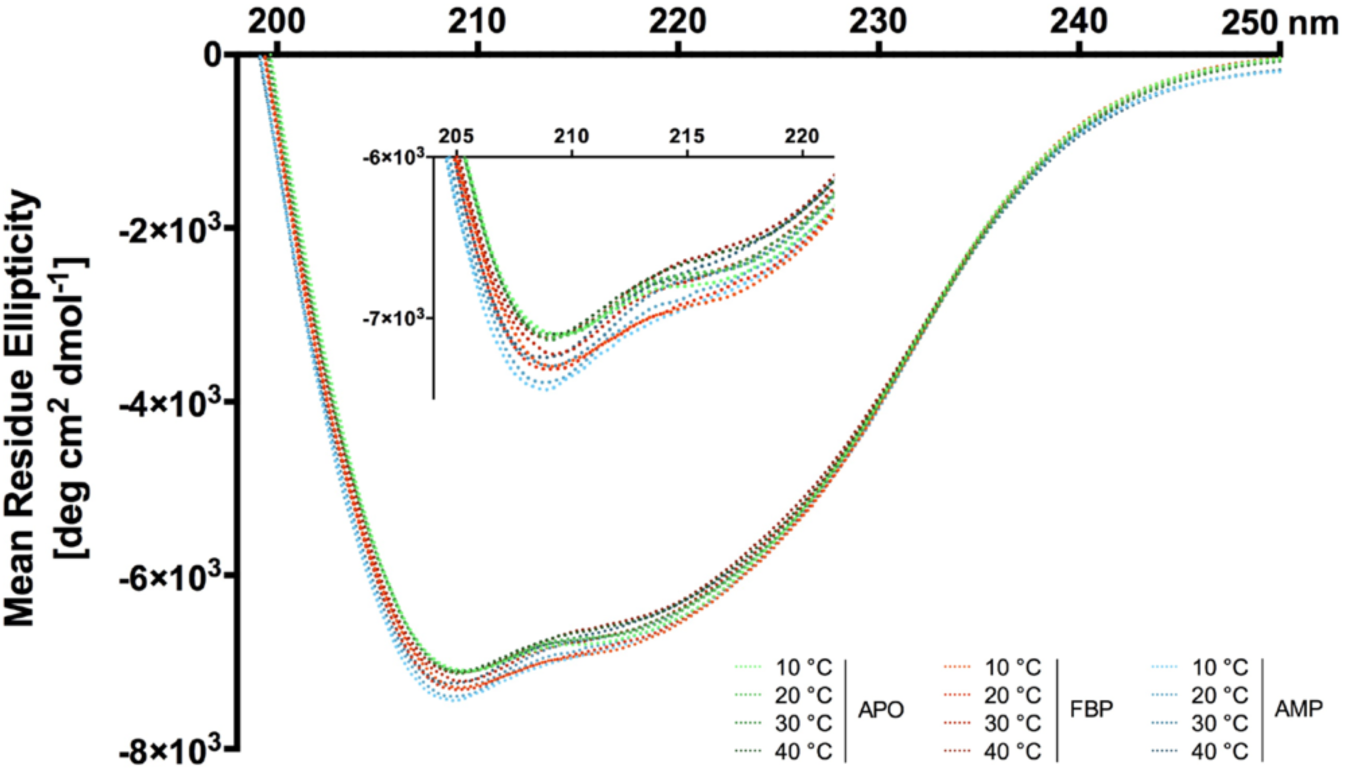
Circular dichroism spectrum of *Ec*AGPase in the absence or presence of regulators at different temperatures. The far-UV CD spectra were acquired at 10, 20, 30 and 40 °C for the *Ec*AGPase in the apo state (green), in the presence of the negative regulator AMP (blue) and in the presence of the activator FBP (red). A close-up view of selected region of the spectra is shown for clarity.

**Figure S12.**
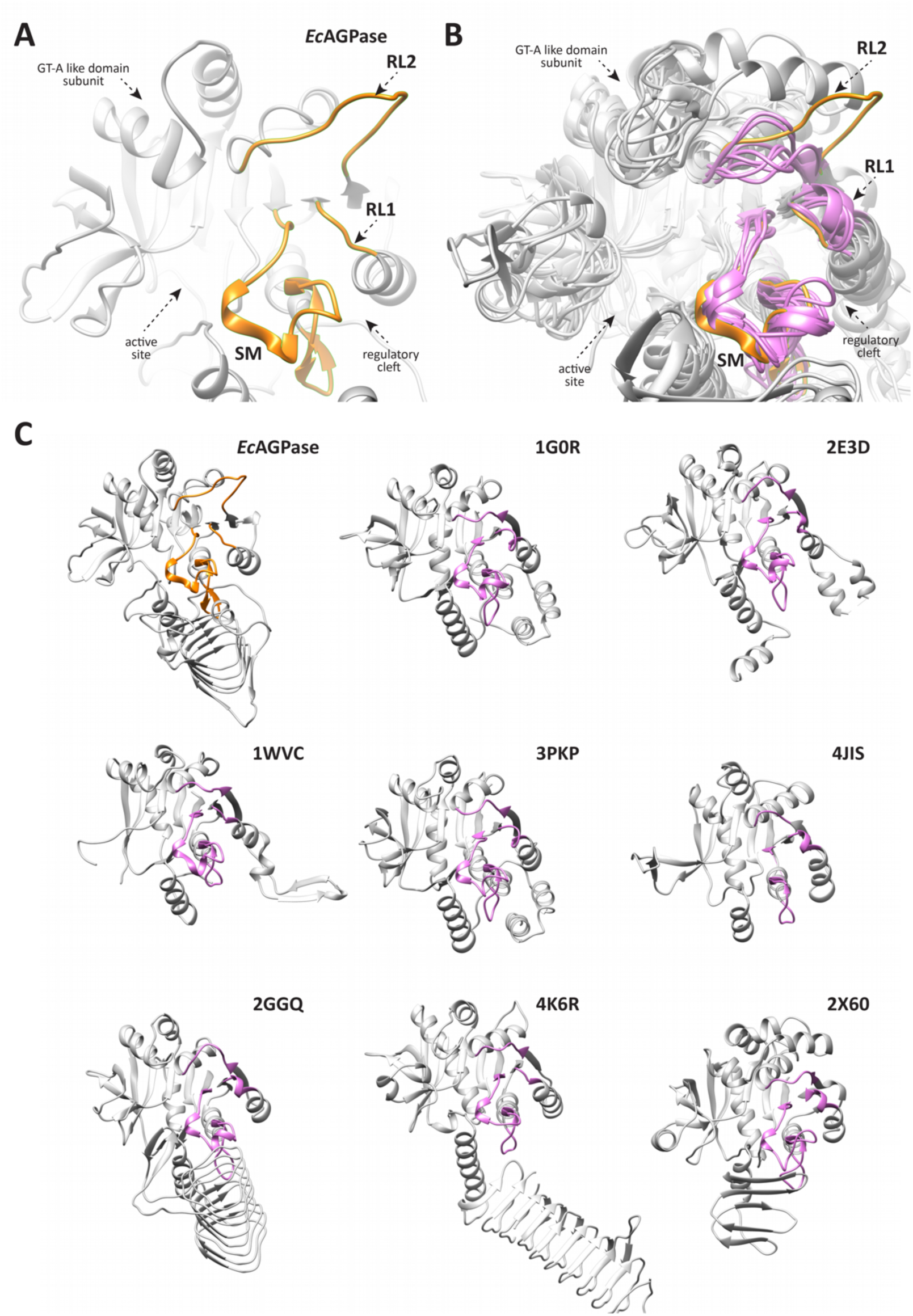
Structural superposition of *Ec*AGPase with other NDPases. CryoEM structure of *Ec*AGPase in complex with AMP (EC: 2.7.7.27); 1G0R: glucose-1-phosphate thymidyltransferase (RmlA) from *P. aeruginosa* in complex with G1P (EC:2.7.7.24); 2E3D: glucose-1-phosphate uridylyltransferase from *E. coli* (EC:2.7.7.9); 1WVC: glucose-1-phosphate cytidylyltransferase from *Salmonella enterica* complexed with cythidine-5’-triphosphate (EC: 2.7.7.33); 3PKP: glucose-1-phosphate thymidyltransferase from *S. enterica* (RmlA-Q83S variant) with 2’deoxy-adenosine triphosphate (EC:2.7.7.24); 4JIS: ribitol 5-phosphate cytidylyltransferase (TarI) from *Bacillus subtilis* (EC:2.7.7.40); 2GGQ: glucose-1-phosphate thymidylyltransferase from *Sulfolobus tokodaii* in complex with TTP (EC:2.7.7.24); 4K6R: GlmU from *Mycobacterium tuberculosis* in complex with ATP (EC: 2.7.7.23); 2X60: GDP-manose pyrophosphorilase from *Thermotoga maritima* in complex with GTP (EC:2.7.7.13).

### SUPPLEMENTARY MATERIALS AND METHODS

#### 3.1. *Ec*AGPase purification

Overnight cultures of 100 ml of *Escherichia coli* BL21 transformed with the plasmid pET22b-*Ec*AGPase containing the full-length *Escherichia coli* K12 *glgC* gene were inoculated in 2L of LB medium supplemented with 100 mg L^-1^ carbenicillin and grow at 37 °C until cultures reached an A600 of 0.8.^16,30^ Cultures were set to 18°C and *Ec*AGPase expression induced by the addition of 1 mM isopropyl-D-1-thio-galactopyranoside (IPTG) and incubated for 18-20 hs. Bacterial cells were harvested by centrifugation at 5,000 g and resuspended in 40 ml of 50mM HEPES pH 8.0, 10% w/v sucrose, 5 mM MgCl_2_ and 0.1 mM EDTA (solution A). The cell suspension was supplemented with protease inhibitors (Pierce protease inhibitor mini tablets, Thermo Scientific), 10 mg L^-1^ of lysozyme (Sigma) and 2 μl of Benzonase (Merck), and incubated for 30 min on an ice bath. Subsequently, the suspension was sonicated and the cell lysate centrifuged for 20 min at 20,000 g. The supernatant was dialyzed two times against solution A, by using a 100 kDa MWCO cellulose dialysis membrane (Spectra/Por, Spectrum). The solution was applied to a 30 ml home packed Q-Sepharose Fast Flow column (GE Healthcare) previously equilibrated with solution A. The column was then washed with solution A until no absorbance at 280 nm was detected. *Ec*AGPase was eluted employing a linear gradient from 100% solution A to 100% solution B (solution A containing NaCl 0.5 M) in 100 ml. Fractions were tested for *Ec*AGPase activity as previously described.^16^ The activite-positive fractions were then pooled. Solid ammonium sulfate was added to the suspension at room temperature to a calculated final concentration of 1.2 M. The resulting suspension was centrifuged at 32,000 g for 20 min, and the collected supernatant applied into a Phenyl HIC PH-814 column (Shodex) previously equilibrated in solution C (solution A containing ammonium sulfate 1.2 M). The enzyme was eluted from the column applying a linear gradient from 100% solution C to 100% solution A in 50 ml at a 1ml min^-1^ flow rate. Fractions with high-purity *Ec*AGPase, tested SDS-page, were pooled and dialyzed against solution A with a 100-kDa molecular overnight at 4°C and stored at 80°C for subsequent experiments.

#### 3.2. CryoEM sample preparation

An initial inspection by negative stain and cryoEM grid-screening and optimization was performed using an in-house Jeol 2200, 200 kV FEG equipped with an UltraScan 4000 CCD camera (Gatan, USA). Frozen aliquots of *Ec*AGPase in solution A where thawed and concentrated to 4 mg ml^-1^. Samples of 50 μl of *Ec*AGPase were injected in an analytical SEC column KW-403-F (Shodex) previously equilibrated with 50 mM Tris-HCl pH 7.5, 100 mM NaCl and 2 mM MgCl_2_, and the protein peak collected in 50 μl fractions for subsequent vitrification. The *Ec*AGPase-AMP sample was obtained by mixing SEC protein fractions at 0.6 mg/ml with an equal volume of a solution of 0.5 mM AMP in 50 mM Tris-HCl, pH 7.5 and 100 mM NaCl. The resulting solution containing *Ec*AGPase 0.3 mg ml^-1^ and 0.25 mM AMP was applied onto copper Quantifoil R 1.2/1.3 grids. Grids were plunge-freeze into liquid ethane using a Vitrobot MkII (FEI) at 10°C and 80 HR% humidity in the chamber (Supplementary Fig. S3). The *Ec*AGPase-FBP complex was obtained mixing equal parts of *Ec*AGPase at 0.6 mg ml^-1^ with 5 mM FBP in 50 mM Tris-HCl pH 7.5 and 100 mM NaCl. The resulting solution containing *Ec*AGPase 0.3 mg ml^-1^ and 2.5 mM FBP was applied onto home-made graphene oxide coated copper Quantifoil R 1.2/1.3 grids using a Vitrobot MkII (FEI) at 10°C and 80 HR% humidity in the chamber (Figure S2).

Home-made graphene oxide grids were made with a smith grid coating trough (LAAD research industries) in which the perforated metal tray was centrally covered with a strip of 1 cm wide Parafilm (Bemis). The device was filled with Milli-Q water until the tray was covered entirely. Grids were immersed in the water and the grids placed on top of the Parafilm, with the carbon side up. A suspension of graphene oxide was made by vortexing 100 μl of graphene oxide at 2 mg ml^-1^ (Sigma) with 100 μl of methanol. The suspension was added drop-by-drop onto the water surface and incubated for 1 min. The liquid in the smith grid coating trough was removed by aspiration with a syringe leaving a drop of water with graphene stayed on top of the grids. The tray with the grids was deposited in a petri dish with filter paper and leave inside a desiccator overnight for next day use.

#### 3.3. CryoEM data collection

Data were collected at the Electron Bio-Imaging Centre (eBIC) at Diamond Light Source, Oxfordshire, UK. The data of both *Ec*AGPase-FBP and *Ec*AGPase-AMP complexes were acquired using the Titan Krios-I (FEI) operated at 300 kV (Cs 2.7 mm). Data was recorded using a Quantum K2 Summit camera (Gatan) in counting mode, recording 40 frames/movie at a final pixel size of 1.047 Å. Automatic data collection was performed using the software EPU 1.9.1 (FEI; Table 1 and Figures S2-S3).

#### 3.4. CryoEM single-particle data processing

*Ec*AGPase-AMP and *Ec*AGPase-FBP complexes data set were processed using the general protocol described in the following lines. Data processing was initiated using the Relion 3.0 suite.^56^ The 40 movie frames of each movie were aligned using MotionCor2 (5×5 patches and Bfactor 150). The CTF of the aligned image was assessed in the non-dose weighted image using CtfFind 4.1 with parameters amplitude contrast 10% and FFT box size of 512 pixels. Image CTF was visually inspected, and images with low quality discarded. Relion reference-free automatic particle picking was performed considering a particle diameter between 95 Å and 110 Å, imposing a minimum inter-particle distance of 90 Å and a low picking threshold (0.05). Extracted particles were extracted using a square box of 200 pixels side. Particles were subjected to several rounds and extensive 2D classification selecting the best classes. An initial first 3D auto-refinement in C1 was performed using a calculated map at 35 Å resolution produced in USCF-Chimera^57,58^ from the *Ec*AGPase tetramer from the crystal structure (pdb code PDB 5L6S). This initial reconstruction was used to re-extract centered particles. Afterward, several steps of 3D classification in C1 were performed to discarding bad classes until models reached resolutions better than 6Å and visual inspection revealed a well-defined tetrameric structure. The selected 3D classes were used to reconstruct a model using Relion 3D auto-refinement protocol in C1, which led to a map resolving better than Å. This reconstruction was used as a starting point to perform one round of Relion CTFrefinement and particle Polishing. Later, polished and CTF-refined particles from Relion were imported into CryoSparc2 and a final 2D classification of 200 classes was performed and the best-defined classes selected (Figures S2B and S3B). Final 3D reconstructions were performed in CryoSparc2 using the homogeneous refinement imposing C1, C2 and D2 symmetry and applying dynamic masking, using the Relion reconstructions low-pass at 30 Å as starting model. Finally, a final step of Non-Uniform refinement for each reconstruction, on the corresponding symmetry, was performed applying a calculated global mask from each reconstruction. The reconstructions obtained from the non-uniform CryoSparc2 protocol are reported for each symmetry. Reported auto sharp-maps were obtained from the CryoSparc2 non-uniform refinements (Figures S2C and S3C). Local resolution was measured in CryoSparc2 LocRes program implementation,^59^ using a threshold for local FSC = 0.5 for local resolution assesment (Figures S2C and S3C). For quality graphs as FSC plots, angular distribution and precision see (Figures S2D-F and S3D-F). For other statistics please see Table 1.

#### 3.5. Model building and refinement

A model of the *Ec*AGPase tetrameter without ligands and trimmed RL1 and RL2 loops from the X-ray crystal structure (pdb code PDB 5L6S), was initially rigid-body fitted inside the cryoEM reconstruction using chimera.^57^ The refinement of the models using Phenix real space_refinement^60,61^ in the sharp maps was performed with alternative rounds of Coot modeling^62^ for and de-novo loop building. Corresponding NCS were enforced for model refinement in the D2 and C2 map reconstructions, meanwhile, automatic NCS determination was used in the C1 reconstruction. The quality of models was checked during refinements using Molprobity and validate-PDB server.^63^ For model statistics and quality details please see Table 1.

#### 3.6. Visualization, structural analysis and bioinformatics

Analysis of the structures and density maps, morphing, sequence alignments, images, and visualization was done using UCSF Chimera. Residue interaction networks analysis was performed using RINalyzer program on Cytoscape. The multiple sequence alignment was performed using UNIPROT online service using Clustal Omega (1.2.4). The aligned AGPase sequences were retrieved from UNIPROT: *Escherichia coli* (P0A6V1), *Triticum aestivum* (P30523), *Arthrobacter sp.* (A0JWV0), *Haemophilus influenzae* (P43796), *Klebsiella pneumoniae* (B5XTQ9), *Enterobacter sp.* (A4WFL3), *Nostoc punctiforme* (B2IUY3), *Rhodobacter sphaeroides* (A3PJX6), *Rhodospirillum rubrum* (Q2RS49), *Rhizobium radiobacter* (P39669), *Solanum tuberosum* (P23509), and *Cronobacter sakazi* (A7MGF4).

#### 3.7. Thermofluor

*Ec*AGPase frozen aliquots were thawed and buffer exchanged in 50 mM Tris-HCl pH 7.5, 100mM NaCl and concentrated to obtain a protein stock of 4mg ml^-1^. The thermofluor assay was performed in a total reaction volume is 25 µl as follows. First, a volume of 20 μl of an assayed mix of ligands in 50 mM Tris-HCl pH 7.5, 100mM NaCl to obtain corresponding final concentrations of either AMP 0.5 mM, FBP 2.5 mM, and/or ADP-glucose 0.5 mM was deposited onto 96-well thin-wall PCR plate (Hardshell, Bio-Rad) in an ice bath. Secondly, 2.5 μl of a 1:100 dilution in water of the probe SYPRO Orange stock (S5692, Sigma) to each well. Finally, 2.5 μl of protein stock or buffer was added corresponding wells to measure protein and controls in triplicates. The reaction plate was covered with transparent adhesive sealer (Microseal, Bio-Rad) and kept on ice bath until measurement. The experiment was performed in a CFX96 qPCR System Thermocycler (Bio-Rad), with the block pre-cooled at 4°C and the lid pre-heated to 98°C. The experiment was run using a ramping temperature from 4°C to 95°C with an increment of 1°C min^-1^ and data measured in the HEX channel. Well-to-well baseline differences between curves were corrected by subtraction their corresponding fluorescence value at the denaturation peak maxima, as considering the fluorescence value at the denaturation peak maxima should be constant to the interaction probe-protein in a given condition. Therefore, the resulting corrected curves with fluorescence value at the denaturation peak maxima equals 0. Corrected curve triplicates were averaged, and the signal from buffers subtracted.

#### 3.8. Isothermal titration calorimetry

To study of binding differences for ATP to *Ec*AGPase in the presence of allosteric regulators was carried out at 25 °C in the presence of 2 mM MgCl_2_ using a MicroCal PEAQ-ITC calorimeter (Malvern Pananalitycal). Specifically, the 20 uM *Ec*AGPase in buffer 100 mM NaCl, 200 mM Tris-HCl pH 7.5 with either AMP 0.1 mM or FBP 2.5 mM was titrated with 14 injections of 2.5 μl of ATP 1 mM in matching buffer with an equilibration time of 300 s/injection. Protein buffers were treated in the same fashion to assess the dilution heat of ATP. Data from protein experiment was subtracted with the control buffers and was fitted imposing a model for four sites sequential binding site using the MicroCal PEAQ-ITC analysis software where: K_D1_=[*Ec*AGPase][ATP]/[*Ec*AGPase-ATP_1_]; K_D2_=[*Ec*AGPase-ATP_1_][ATP]/[*Ec*AGPase-ATP_2_]; K_D3_=[*Ec*AGPase-ATP_2_][ATP]/[*Ec*AGPase-ATP_3_]; K_D4_=[*Ec*AGPase-ATP_3_][ATP]/[*Ec*AGPase-ATP4].

#### 3.9. Far-UV Circular Dichroism spectra and thermal unfolding

*Ec*AGPase frozen aliquots in solution A thawed and concentrated to 4mg/ml. These samples were buffer exchanged by size-exclusion chromatography (Superdex 200 10/300 GL, GE) into buffer 10 mM Tris pH 7.5 100 mM NaF (CD buffer). Protein fractions of the single peak corresponding to the EcAGPase tetramer were pooled to a final concentration of 0.6 mg/ml. Samples were prepared by mixing equal parts of 0.6 mg/ml protein solution with CD buffer with either 0.2 mM AMP, 5 mM FBP, or without regulators. The CD spectra in the far UV region (250-190 nm) were measured utilizing a J-815 CD spectrometer (Jasco), by using a 1 mm path length High Precision Cell cuvette (Hellma). To explore the ligand effects in a range of physiological temperatures, 20 accumulations of each spectrum were recorded for each sample at 10, 20, 30 and 40 °C. Spectra were normalized by subtracting the buffer spectra from the raw data. Data were converted from mdeg to Mean Residue Ellipticity [*θ*]_MRW,_*λ* according to the formula “[*θ*]_MRW,_*λ* = *θ*.MRW/10.d.c”, where *θ* is the observed ellipticity (degrees), MRW is the molecular mass divided by N-1, with N as the number of amino acids, d is the optical path (cm), and c is the concentration in g/ml.

#### 3.10. Qualitative meta-analysis of *Ec*AGPAse single point mutants

To simplify the examination of reported kinetic properties of *Ec*AGPase single-point mutants, we approach the meta-analysis only for single point mutations of residues at the active site and the regulatory cleft to alanine residues (Table 2).^32,37,38^ In addition, to set the basis for the comparison across different reports, we standardized mutant values as a ratio against the corresponding wild-type value reported in each individual study, i.e. Vmax-ratio = mutantVmax/wild-typeVmax. Finally, for the benefit of a qualitative characterization of these mutants, we perform an *ad-hoc* classification into three groups with marked differences: (i) mutants with parameter between the half and the double of the values displayed by the wild-type enzyme (0.5> parameter mutant/wild-type < 2.0, symbolized as “∼” in Table 2), (ii) mutants with high ratio (parameter mutant/wild-type > 2.0, symbolized as “↑” in Table 2), and (iii) mutants with low ratio (parameter mutant/wild-type < 0.5, symbolized as “↓” in Table 2).

